# Recurrent chromosomal translocations in sarcomas create a mega-complex that mislocalizes NuA4/TIP60 to Polycomb target loci

**DOI:** 10.1101/2021.03.26.436670

**Authors:** Deepthi Sudarshan, Nikita Avvakumov, Marie-Eve Lalonde, Nader Alerasool, Charles Joly-Beauparlant, Karine Jacquet, Amel Mameri, Jean-Philippe Lambert, Justine Rousseau, Catherine Lachance, Eric Paquet, Lara Herrmann, Samarth Thonta Setty, Jeremy Loehr, Marcus Q. Bernardini, Marjan Rouzbahman, Anne-Claude Gingras, Benoit Coulombe, Arnaud Droit, Mikko Taipale, Yannick Doyon, Jacques Côté

## Abstract

Chromosomal translocations frequently promote carcinogenesis by producing gain-of-function fusion proteins. Recent studies have identified highly recurrent chromosomal translocations in patients with Endometrial Stromal Sarcomas (ESS) and Ossifying FibroMyxoid Tumors (OFMT) leading to an in-frame fusion of PHF1 (PCL1) to six different subunits of the NuA4/TIP60 complex. While NuA4/TIP60 is a co-activator that acetylates chromatin and loads the H2A.Z histone variant, PHF1 is part of the Polycomb repressive complex 2 (PRC2) linked to transcriptional repression of key developmental genes through methylation of histone H3 on lysine 27. In this study, we characterize the fusion protein produced by the *EPC1*-*PHF1* translocation. The chimeric protein assembles a mega-complex harboring both NuA4/TIP60 and PRC2 activities and leads to mislocalization of chromatin marks in the genome, in particular over an entire topologically-associating domain including part of the *HOXD* cluster. This is linked to aberrant gene expression, most notably increased expression of PRC2 target genes. Furthermore, we show that JAZF1, implicated with a PRC2 component in the most frequent translocation in ESS, *JAZF1-SUZ12*, is a potent transcription activator that physically associates with NuA4/TIP60, its fusion creating similar outcomes as *EPC1-PHF1*. Importantly, the specific increased expression of PRC2 targets/*HOX* genes was also confirmed with ESS patient samples. Altogether, these results indicate that most chromosomal translocations linked to these sarcomas employ the same molecular oncogenic mechanism through a physical merge of NuA4/TIP60 and PRC2 complexes leading to mislocalization of histone marks and aberrant polycomb target gene expression.

## INTRODUCTION

ATP-dependent remodelers and histone modifiers are key regulators of the structure and function of chromatin, essential for genome expression and stability, cell proliferation, development and response to environmental cues. Post-translational modifications of specific residues on histones are also part of the epigenetic mechanisms ensuring gene expression memory during cell divisions (Zentner and Henikoff 2013). Histone modifying enzymes are often part of large multisubunit protein complexes. They are highly conserved and composed of various combinations of subunits like readers, writers, and erasers of histone marks, histone chaperones, and chromatin remodelers. The combination of different modules ensures specific localization, histone target selection, enables epigenetic crosstalk and context-specific activity. The strictly coordinated activities of chromatin modifying complexes ensure the proper functioning of the cell (Lalonde et al. 2014). Disruption of chromatin marks and their regulators can lead to various pathologies including cancer (Shen and Laird 2013).

Recurrent chromosomal translocations producing oncogenic fusion proteins are common in many cancers and particularly prevalent in hematopoietic malignancies and sarcomas. These fusions frequently involve chromatin and transcription regulators and, in many cases, are thought to be the primary drivers of cancer. In recent years, disruption of chromatin dynamics has emerged as a consistent oncogenic mechanism used by these fusion proteins (reviewed in (Brien et al. 2019)). Sarcomas are rare mesenchymal tissue cancers with distinct molecular profiles. 1/3rd of all sarcomas harbor chromosomal translocations with otherwise normal karyotypes, showing robust clustering of gene expression profiles compared to normal tissues. Recurrent translocations are found in Endometrial Stromal Sarcomas (ESS) and Ossifying FibroMyxoid Tumors (OFMT) that potentially fuse subunits of distinct chromatin modifying complexes with opposite functions in gene regulation, namely the NuA4/TIP60 histone acetyltransferase (HAT) complex and Polycomb Repressive complexes (Figure 1A). Among the recurring chromosomal translocations that characterize ESS, genes for five different subunits of the NuA4/TIP60 complex (EPC1/2, MBTD1, MEAF6, BRD8) are repeatedly found fused to genes for different PRC2 components (PHF1/SUZ12/EZHIP) (Ferreira et al. 2018; Hoang et al. 2018; Momeni-Boroujeni et al. 2021). Interestingly, other soft tissue sarcomas such as OFMT also harbor EPC1-PHF1, EP400-PHF1 and MEAF6-PHF1 fusions. OFMTs are rare cancers of uncertain cellular origin, and ∼50-85% of cases show PHF1 translocations, with EP400-PHF1 being the most frequent (40%) (Schneider et al. 2016). JAZF1-PHF1 was reported in a case of Cardiac Ossifying sarcoma and fusions of NuA4/TIP60 subunits with the PRC1.1 component BCOR have also been reported in ESS, demonstrating that these types of fusion events represent a critical oncogenic mechanism in mesenchymal cancers (Schoolmeester et al. 2013) (Figure 1A).

**Figure 1.**
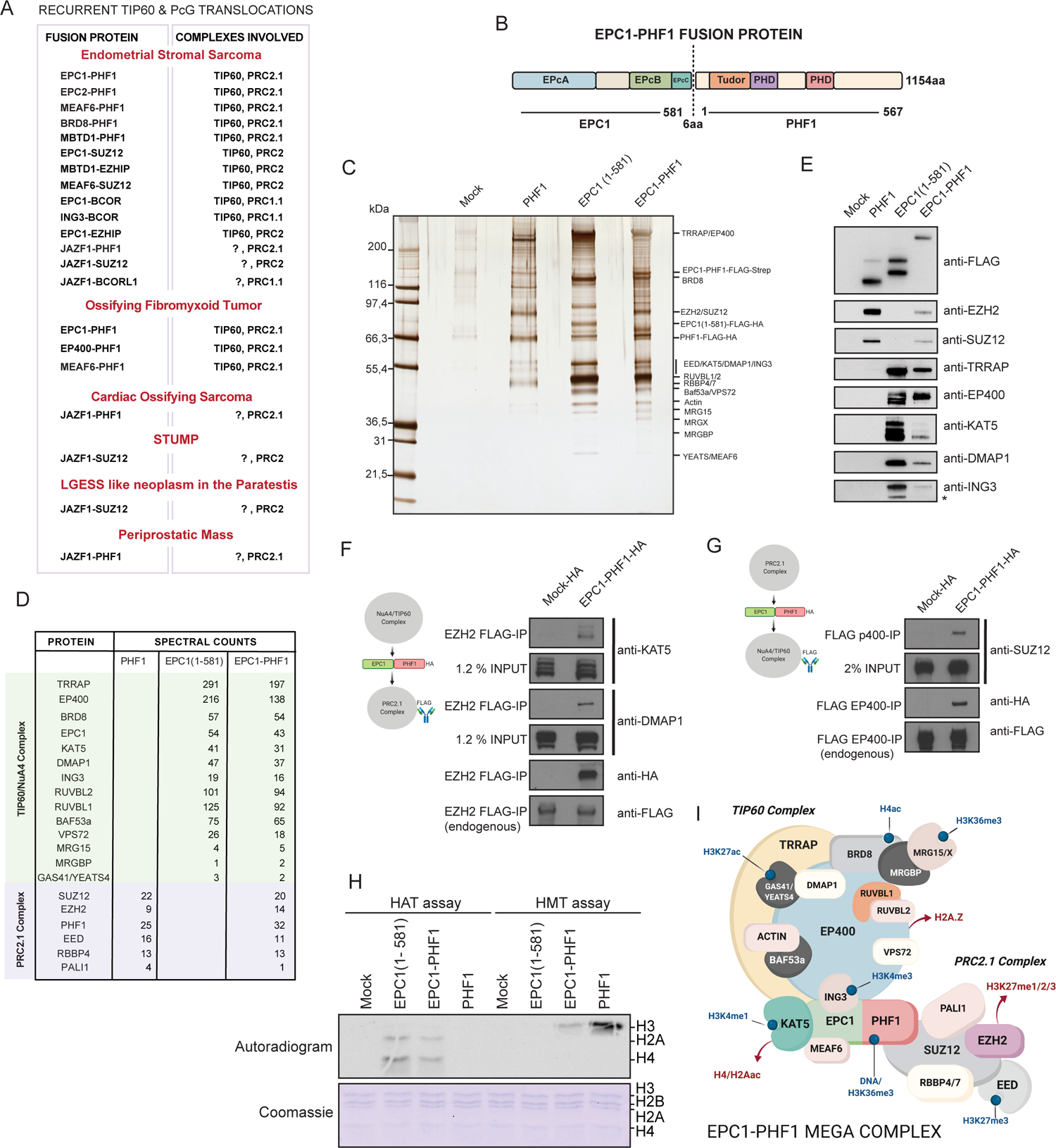
The EPC1-PHF1 fusion protein assembles a mega-complex merging NuA4/TIP60 and PRC2 complexes, and their activities. **(A)** Table summarizing recurrent translocations found in soft-tissue sarcomas that fuse NuA4/TIP60 complex proteins and Polycomb group proteins. **(B)** Schematic representation of the EPC1-PHF1 fusion protein. The numbers indicated are amino acids, protein domains retained in the fusion are indicated. **(C)** Silver stained SDS-PAGE showing affinity purified complexes. Labels on the right side are proteins that were identified based on western blotting and predicted molecular weights. **(D)** Western blots of selected NuA4/TIP60 and PRC2 complex subunits on the affinity purified complexes shown in (C). **(E)** Mass Spectrometry analysis of the affinity purified complexes shown in (C). See also Figure S2A-B. **(F-G)** Affinity Purification of endogenous PRC2 complex (EZH2 subunit) (F) and NuA4/TIP60 (EP400 subunit) (G) and confirming formation of a merged mega-complex by EPC1-PHF1. **(H)** *In vitro* Histone Acetyltransferase (HAT) assay and Histone Methyltransferase (HMT) assay. KAT5 purified through EPC1(1-581) as well as EPC1-PHF1 fusion show HAT activity towards H2A and H4 on chromatin (Upper panel, Lane 2 and 3). EZH2 purified through PHF1 as well as EPC1-PHF1 fusion show HMT activity towards H3 on chromatin (Upper panel, Lane 7 and 8). Coomassie stained SDS-PAGE gel is the loading control for nucleosomal histones. See also Figures S2C and D. **(I)** Schematic representation of the chimeric mega-complex assembled by the EPC1-PHF1 fusion protein.

NuA4/TIP60 is an evolutionarily conserved and multi-functional MYST-family HAT complex with 17 distinct subunits (Steunou et al. 2014; Judes et al. 2015; Jacquet et al. 2016; Sheikh and Akhtar 2019). The catalytic subunit KAT5/Tip60 acetylates histones H4 and H2A, as well as variants H2A.Z and H2A.X in the context of chromatin. In addition, the ATP-dependent remodeler subunit EP400 allows the exchange of canonical histone H2A in chromatin with variant histone H2A.Z (Billon and Cote 2013; Pradhan et al. 2016). There are also multiple subunits with different reader domains that allow context-dependent site-specific activity of the complex and epigenetic crosstalk (Steunou et al. 2014; Jacquet et al. 2016). The 836aa EPC1 protein is a non-catalytic scaffolding subunit of the TIP60 complex. The conserved N-terminal EPcA domain (aa1-280) interacts with KAT5/Tip60, MEAF6, and ING3 thereby forming the human Piccolo NuA4 complex, enabling binding and acetylation of chromatin substrates (Boudreault et al. 2003; Doyon et al. 2004; Selleck et al. 2005; Lalonde et al. 2013; Xu et al. 2016). The C-terminal part of EPC1 is thought to associate with the rest of the NuA4/TIP60 complex based on data from the yeast NuA4 complex (Boudreault et al. 2003; Setiaputra et al. 2018; Wang et al. 2018) (Figure 1B). The NuA4/TIP60 complex is an important transcriptional co-activator that can be recruited to gene regulatory elements by several transcription factors. Through acetylation of histones as well as non-histone substrates, it activates a multitude of gene expression programs, including: proliferation, stress response, apoptosis, differentiation and stem cell identity (Steunou et al. 2014; Judes et al. 2015; Sheikh and Akhtar 2019). The NuA4/TIP60 complex is also a key player in the response to DNA damage, assisting repair pathway choice as well as the repair process itself ((Jacquet et al. 2016; Cheng et al. 2018) and references therein). All these functions are essential for cellular homeostasis, hence NuA4/TIP60 subunits are often mutated or deregulated in cancers and Tip60/KAT5 itself is a haplo-insufficient tumor suppressor (Gorrini et al. 2007; Judes et al. 2015; Sheikh and Akhtar 2019).

Polycomb Group proteins (PcG) are evolutionarily conserved proteins involved in development and transcriptional regulation, originally linked to *HOX* gene repression during Drosophila development (Schuettengruber et al. 2017). They are key regulators of mammalian development, differentiation, and cell fate decisions. PcG proteins assemble into multisubunit complexes, major ones being Polycomb Repressive Complexes 1 and 2 (PRC1 and PRC2). These complexes co-operate with each other to create repressive chromatin regions through histone modifications and chromatin organization. While PRC1 catalyzes H2AK119 mono-ubiquitination and chromatin compaction, PRC2 catalyzes H3K27 methylation. The PRC2 complex is composed of the core components EZH2, SUZ12, EED, and RBBP4/7. Through a read-write mechanism, it can deposit H3K27me3 over large chromatin regions such as the *HOX* clusters. Other associated factors function to stabilize the PRC2 complex on chromatin and modulate its activity, defining distinct variant complexes PRC2.1 and PRC2.2. PRC2.1 contains one of the Polycomb-like (PCL) proteins, PHF1, MTF2 or PHF19, as well as either PALI1/2 or EPOP (Laugesen et al. 2019; Loubiere et al. 2019). The 567aa PHF1 protein binds unmethylated CpG islands particularly at long linker DNA regions, stabilizes the binding of PRC2.1 complex on chromatin and increases its activity (Cao et al. 2008; Sarma et al. 2008; Choi et al. 2017; Li et al. 2017). PHF1 is also known to bind the H3K36me3 mark through its Tudor domain (Musselman et al. 2012; Cai et al. 2013), restricting the catalytic activity of EZH2.

Many groups are working on the classification of ESS using recurrent fusions as molecular markers. However, there is currently little molecular understanding about the consequences of these recurrent translocations and their primary role in tumorigenesis. To address this, we used biochemical and genomic approaches to study the product of a recurrent translocation found in ESS and OFMTs that fuses the EPC1 subunit of the NuA4/TIP60 complex to the PHF1 subunit of the PRC2.1 complex (Figure 1B)(Micci et al. 2006; Antonescu et al. 2014). We investigated the molecular impact of the EPC1-PHF1 fusion protein by generating isogenic cell lines, used affinity purification followed by mass spectrometry to identify interactors, associated activities and analyzed chromatin occupancy of the fusion protein. We also analyzed the effect of the fusion protein on global chromatin dynamics while correlating differential gene expression.

## RESULTS

To construct the fusion *EPC1-PHF1* gene (Figure 1B), we used a portion of the chimeric gene recovered by RT-PCR from total RNA isolated from surgically removed endometrial stromal sarcoma (a kind gift from Francesca Micci’s group) (Micci et al. 2006). As necessary controls, we also subcloned full-length PHF1 and EPC1, as well as a portion of EPC1 corresponding to the fragment found in the fusion, referred to as EPC1(1-581). We confirmed the expression of all constructs and then tested whether the EPC1-PHF1 fusion protein can act like an oncogenic driver by performing Colony Formation Assay in HEK293T cells (Figure S1A-B). In contrast to full-length and truncated EPC1 which strongly inhibit growth, we observed that expression of EPC1-PHF1 leads to a greater number of colonies compared to controls (Figure S1B), supporting the idea that this gene fusion may be a driver event, giving a growth advantage to the expressing cells.

### EPC1-PHF1 forms a mega-complex merging NuA4/TIP60 and PRC2.1

To isolate the native complex(es) formed by the EPC1-PHF1 fusion protein, we generated isogenic K562 cell lines by targeted integration of C-terminally TAP-tagged cDNAs into the *AAVS1* safe harbor genomic locus (Dalvai et al. 2015). To circumvent issues related to protein overexpression, we used the moderately active *PGK1* promoter which was shown to achieve expression of NuA4/TIP60 and PRC2 subunits within 2.5-fold of the native levels (EP400, EPC1, MBTD1 and EZH2) (Dalvai et al. 2015). Our previous studies have demonstrated that this system enables purification of stable and stoichiometric protein complexes (Dalvai et al. 2015; Doyon and Cote 2016; Jacquet et al. 2016). We chose the K562 cell line as it is an ENCODE tier-1 cell line and can grow in suspension culture to high cell densities, ideal for biochemical and genomic studies.

Nuclear extracts from K562 cell lines expressing the fusion protein, individual fusion partners or the empty tag (Mock, C-terminal 3xFlag-2xHA-2A-puromycin tag) were allowed to bind to anti-FLAG resin and the bound material was eluted with 3xFlag peptides. An alternate K562 cell line expressing EPC1-PHF1 with a C-terminal 3xFLAG-2xStrep tag was also used to improve yield and purity by binding the FLAG eluted fraction to Streptactin beads followed by elution with biotin. SDS-PAGE and silver-staining identified the components of the purified complexes (Figure 1C). Mass spectrometry analysis identified all expected components of NuA4/TIP60 and PRC2 complexes (Figure 1D, S2A-B). We further confirmed the complex subunits by western blotting with NuA4/TIP60 and PRC2 specific antibodies (Figure 1E). The PHF1 fraction contained the core subunits EZH2, EED and SUZ12 of the PRC2 complex, as well as RBBP4 and the PRC2.1 specific PALI1. The EPC1(1-581) fraction contained all the subunits of NuA4/TIP60 complex except MBDT1, which was expected since it associates through an interaction with EPC1 C-terminus (aa644-672) (Jacquet et al. 2016; Zhang et al. 2020). Strikingly, the EPC1-PHF1 fraction contained subunits of both the TIP60 and PRC2.1 complexes. These results indicate that the EPC1-PHF1 fusion protein efficiently associates with both NuA4/TIP60 and PRC2.1. This was also supported by data in HEK293T cells (Figure S1C).

To determine if the fusion protein’s association with the two complexes occur independently of each other or could occur simultaneously, we integrated 3xHA-tagged EPC1-PHF1 at the *AAVS1* locus in K562 cell lines in which TALEN/CRISPR was used to introduce a 3xFLAG tag at endogenous *EZH2* or *EP400* genes, respectively (Dalvai et al. 2015). This allowed us to purify endogenous PRC2 and NuA4/TIP60 complexes and verify if the fusion protein can physically bridge the two complexes. As shown in Figure 1F-G, it is clear that expression of the EPC1-PHF1 fusion leads to the association of the PRC2 complex with NuA4/TIP60 and vice versa, demonstrating the formation of a mega-complex.

We then performed *in vitro* histone acetyltransferase (HAT) and histone methyltransferase (HMT) assays with purified fractions on native chromatin as substrate. We observed robust HAT activity towards histone H4 and H2A in both EPC1(1-581) and EPC1-PHF1 complexes, as expected for NuA4/TIP60 (Figure 1H, S2C). Simultaneously, we detected HMT activity towards histone H3 in PHF1 and EPC1-PHF1 complexes, as expected for PRC2 (Figure 1H, S2D). This demonstrates that the merged assembly maintains the enzymatic activities of both original complexes. Altogether, these results indicate that expression of the EPC1-PHF1 fusion protein in cells leads to the formation of a hybrid TIP60-PRC2 mega-complex that harbours opposite functions in term of chromatin modifications and impact on gene regulation (Figure 1I). It is important to note that we did not detect an effect of expressing the fusion protein or its partners on the bulk level of H3K27me3 (Figure S2E), in contrast to what has been previously proposed for the JAZF1-SUZ12 fusion (Ma et al. 2017).

### The EPC1-PHF1 fusion complex is enriched at genomic loci bound by TIP60 and PRC2.1

To determine the genomic locations where the EPC1-PHF1 mega-complex may act, we performed anti-FLAG chromatin immunoprecipitation coupled to high-throughput sequencing (ChIP-seq) using the isogenic cell lines. We analyzed binding regions (significant peaks) of EPC1-PHF1, EPC1(1-581) and PHF1 in comparison to the empty vector cell line. While over 20 000 bound regions were identified for each TIP60(EPC1) and PRC2.1(PHF1) complexes, only 2 888 were called for the fusion, likely due to its lower expression (Figure 2A-C). Nevertheless, we focused on these locations bound by EPC1-PHF1 and determined overlaps with the regions bound by TIP60, PRC2.1 or unique to the fusion. While most of the sites bound by the fusion overlap with TIP60-bound regions, a significant number also overlaps with PRC2.1 sites and with regions bound by both PRC2.1 and TIP60 (Figure 2A-C, Table S1). Only a small number of sites unique to the fusion is detected. These results indicate that the EPC1-PHF1 fusion protein is targeted to genomic locations normally bound by TIP60, PRC2.1 or both.

**Figure 2.**
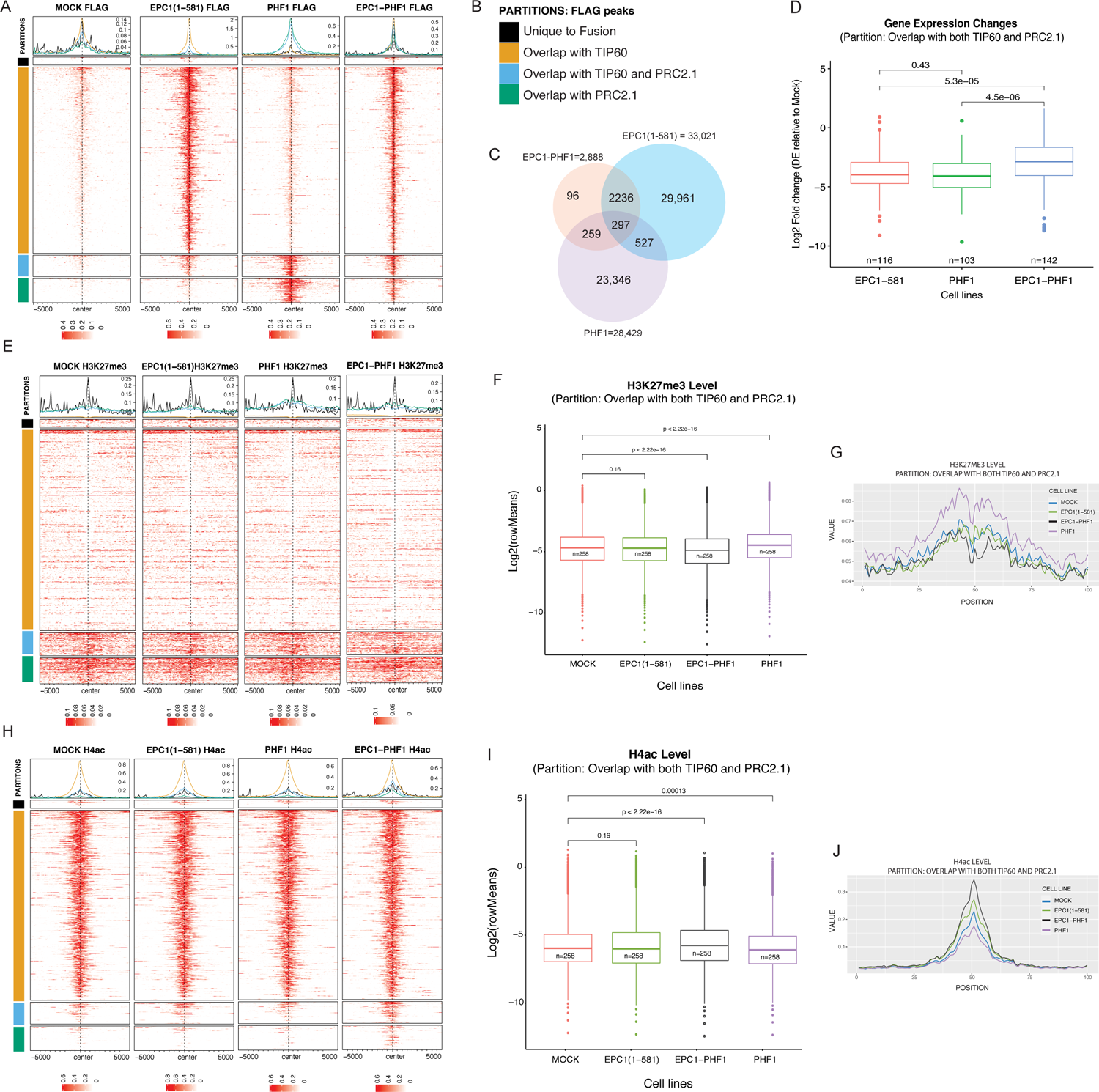
EPC1-PHF1 fusion complex induces global changes in histone marks and gene expression. **(A)** Enrichment of FLAG ChIP-seq signal in K562 cell lines. The heatmaps show regions bound by EPC1-PHF1, partitioned based on overlap with EPC1(1-581) and/or PHF1 FLAG peaks. **(B)** Partitions used for heatmaps in Figure 2A, 2E and 2H. EPC1 (1-581) = TIP60, PHF1 = PRC2.1 **(C)** Venn diagram showing the number FLAG ChIP-seq peaks and overlap between EPC1(1-581), PHF1, and EPC1-PHF1 expressing K562 cell lines. **(D)** Gene expression changes at genes bound by EPC1-PHF1 in a particular partition (EPC1-PHF1 bound regions and overlap with TIP60 and PRC2.1) in the different isogenic cell lines versus mock control cell line. See Figure S4B-D for other partitions. p-value calculated by Wilcoxon testing. **(E)** Enrichment of H3K27me3 ChIP-seq signal in K562 cell lines at regions bound by EPC1-PHF1, partitioned based on overlap with EPC1(1-581) and/or PHF1 FLAG peaks. **(F)** Boxplots showing H3K27me3 enrichment level in K562 cell lines in a particular partition (EPC1-PHF1 bound regions and overlap with TIP60 and PRC2.1). See Figure S3 for other partitions. Statistics were computed using bins; n= number of regions analyzed where 1 region is 100 bins. p-value calculated by Wilcoxon testing. **(G)** Density plot of H3K27me3 enrichment in K562 cell lines in a particular partition (EPC1-PHF1 bound regions and overlap with TIP60 and PRC2.1). See Figure S3 for other partitions. **(H)** Enrichment of H4-penta-acetyl ChIP-seq signal at regions bound by EPC1-PHF1, partitioned based on overlap with EPC1(1-581) and/or PHF1 FLAG peaks. **(I)** Boxplots showing H4ac enrichment level in K562 cell lines in a particular partition (EPC1-PHF1 bound regions and overlap with TIP60 and PRC2.1). See Figure S3 for other partitions. Statistics were computed using bins; n= number of regions analyzed where 1 region is 100 bins. p-value calculated by Wilcoxon testing **(J)** Density plot of H4ac enrichment in K562 cell in a particular partition (EPC1-PHF1 bound regions and overlap with TIP60 and PRC2.1). See Figure S3 for other partitions.

### Effect of EPC1-PHF1 on the chromatin and transcriptional landscape

Two major possible effects are expected from the presence of the fusion mega-complex at specific genomic loci: 1-appearance of chromatin acetylation where PRC2.1 is normally bound on silenced regions of the genome; 2-appearance of H3K27 methylation where TIP60 is normally bound on expressed regions of the genome. We also speculated that the fusion protein may mislocalize TIP60-mediated H4 acetylation to poised bivalent chromatin regions, pushing them towards transcriptional activation. To test this, we performed H4 Penta-acetyl (RRID:AB_310310) and H3K27me3 (RRID:AB_27932460) ChIP-sequencing in our isogenic cell lines. Analysis of these histone marks on the four partitions of the sites bound by EPC1-PHF1 uncovered interesting significant changes. On regions that overlap with both TIP60 and PRC2.1, significant increase of H4 acetylation and decrease of H3K27 methylation are seen on cells expressing the fusion (Figure 2E-J, Table S1). An increase of H4 acetylation is also detected on sites overlapping with PRC2.1 alone, while a decrease of H3K27me3 is less clear although obvious when only comparing to PHF1-expressing cells (Figure S3A-D). EPC1-PHF1 bound sites overlapping with TIP60 alone don’t show striking difference in term of acetylation while the drop of H3K27me3 is difficult to judge because of the very low starting level of this mark at these locations (Figure S3E-H). For the sites unique to the fusion, the low number makes it difficult to judge but there seems to be an increase of acetylation (Figure S3I-L). Altogether, these results indicate that binding of the EPC1-PHF1 fusion alters the epigenome most strikingly by increasing H4 acetylation and decreasing H3K27 methylation at regions bound by both TIP60 and PRC2.1, as well as increasing acetylation at regions bound by PRC2.1 alone.

To understand the effect of EPC1-PHF1-induced changes in local histone modifications on transcription, we performed gene expression analysis of the isogenic cell lines (Figure S4A, Table S1). Again focusing on the partition of the sites bound by EPC1-PHF1, genes neighboring regions that overlap with both TIP60 and PRC2.1 show significantly increased transcription in cells expressing the fusion (Figure 2D). Changes in neighboring gene expression are less striking at the other partitions of EPC1-PHF1-bound sites, although those overlapping with PRC2.1 also show an increase (Figure S4B-D). Overall, the results indicate that EPC1-PHF1-induced increase in H4 acetylation correlate with increased local gene expression. In parallel, we do not clearly detect locations bound by the fusion that show increased H3K27me3 and decreased transcription. On the contrary, the H3K27me3 mark deposited by PRC2 tends to decrease on regions with overlapping binding of TIP60 and PRC2.1, as local transcription increases. Interestingly, these regions could be linked to bivalent chromatin as TIP60 is normally enriched on H3K4me3-containing regions.

Since the increase of H4 acetylation signals at specific EPC1-PHF1-bound regions is predominant, this suggests that subverting NuA4/TIP60 activity to new targets by the fusion protein is the primary outcome. We found that the *HOXD* gene cluster, in particular, demonstrates this effect in a striking fashion. Representative ChIP-seq tracks of FLAG, H4 acetylation and H3K27me3 at the posterior *HOXD* genes in the isogenic cells are shown in Figure 3A. Binding of EPC1(1-581) alone is detected on two precise locations coinciding with small pre-existing H4 acetylation islands at the end of the *EVX2* gene and between *HOXD12* and *HOXD13*. In contrast, PHF1 associates with the entire region, as expected from the H3K27me3 signal detected throughout the same region. Interestingly, while the H3K27me3 may be increased by the expression of exogenous PHF1, the two small H4 acetylation islands persist. Most strikingly, the EPC1-PHF1 cell line, while expressing much less fusion protein compared to EPC1(1-581) and PHF1 (Figure S5A), clearly demonstrates appearance and mislocalization of H4 acetylation over a relatively large region (∼25kb highlighted in yellow). Importantly, we can also observe a decrease in the levels of H3K27me3 in this same region, compared to the isogenic cell lines and also compared internally to the neighboring region (starting from the *HOXD12* gene). This local increase in H4 acetylation correlates with specific increase in expression of the *EVX2* and *HOXD13* genes located within the region (Figure 3B). The Flag signals reflecting EPC1-PHF1 binding on this region were too low to be called a peak but nevertheless show higher mean reads over the increased acetylation peaks compared to mock cells (Figure S5G). To clearly demonstrate that EPC1-PHF1 is binding at these locations, we performed a CUT and RUN-seq experiment with an isogenic cell line expressing an HA-tagged fusion (Table S1). With this more sensitive approach with lower background, specific EPC1-PHF1 peaks are detected at the locations where new H4 acetylation appears (Figure 3C). To validate these observations, we performed ChIP-qPCR with primers to four sites of *de novo* H4 acetylation (*HOXD13*-A, B, C, and D highlighted in Figure 3A). Correcting for nucleosome occupancy with total H2B signal, we could confirm the specific increase in H4 acetylation at these locations in the cell line expressing EPC1-PHF1 (Figure 3D).

**Figure 3.**
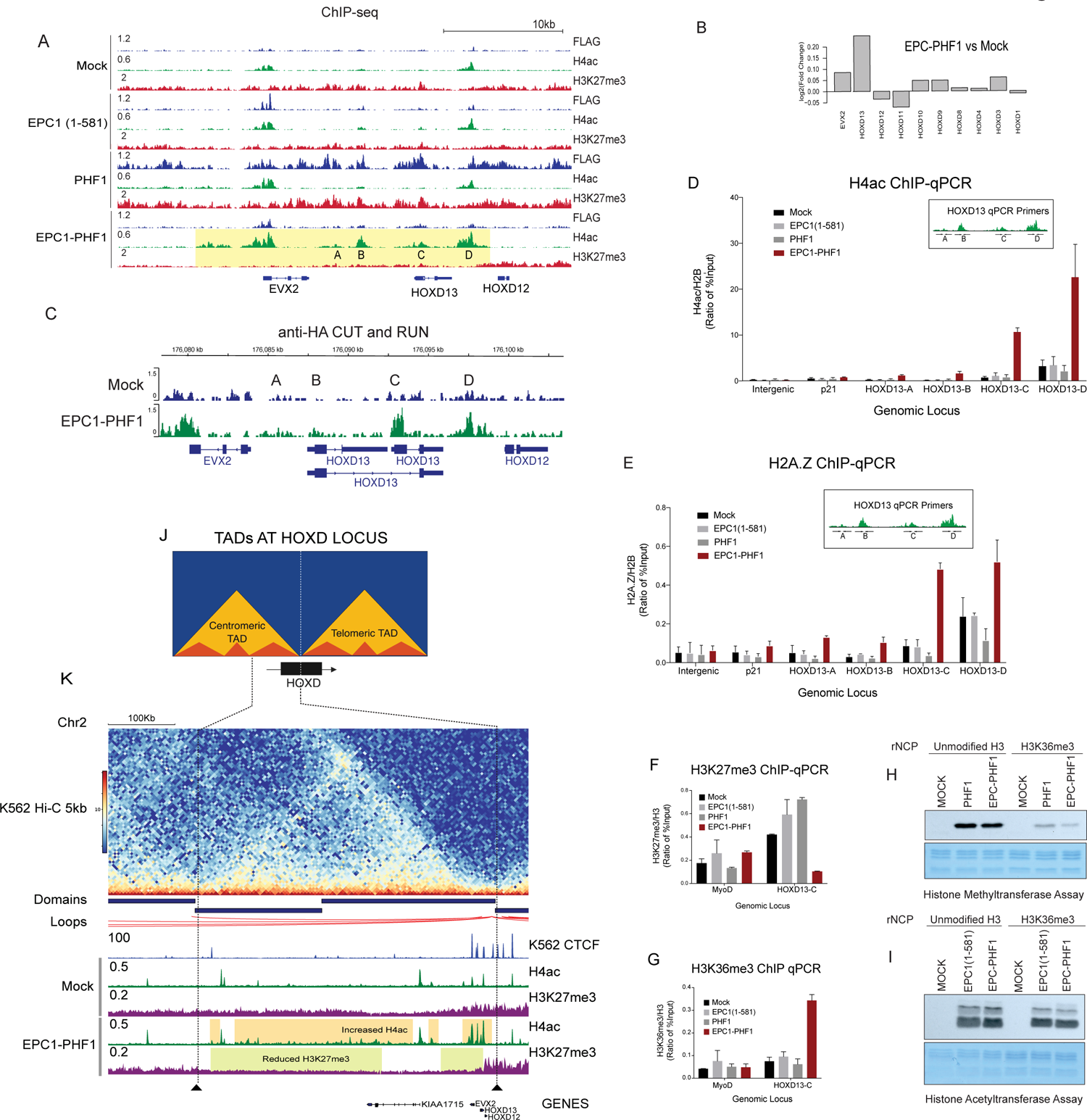
EPC1-PHF1 induces changes in the chromatin landscape at the *HOX* gene clusters. **(A)** Representative ChIP-seq peaks at the *HOXD* gene cluster, highlighted region in yellow shows H4 acetylation mislocalization (at regions labelled *HOXD13*-A, B, C, D) and reduction in H3K27me3 in EPC1-PHF1 expressing K562 cell line. See Figure S6 for similar effects at the other *HOX* clusters and Figure S5D for similar effects in other genes. **(B)** Gene expression changes at the *HOXD* cluster in K562 cells expressing EPC1-PHF1 compared to cells expressing tag only (Mock). See Figure S4A for genome wide gene expression changes. **(C)** Confirmation of EPC1-PHF1 localization at *HOXD13* by anti-HA CUT and RUN sequencing. Representative peaks at the *HOXD13* gene. **(D)** Mislocalization of H4 acetylation determined by ChIP-qPCR. EPC1-PHF1 expressing K562 cell line (Bar graph in maroon color) shows increased H4 acetylation compared to control cell lines at *HOXD13*-A, B, C, D regions, validating the results of ChIP-seq in (A). (Values are a ratio of %Input of H4ac and H2B) (n=2, error bars are range of the values). Intergenic (chr12: 65,815,182-65,815,318) and p21 promoter are negative controls. **(E)** Mislocalization of variant histone H2A.Z determined by ChIP-qPCR. EPC1-PHF1 expressing K562 cell line (Bar graph in maroon color) show increased H2A.Z occupancy compared to control cell lines at regions of H4 acetylation mislocalization (*HOXD13*-A, B, C, D) shown in (A). (Values are a ratio of %Input of H4ac and H2B) (n=2, error bars are range of the values). Intergenic (chr12: 65,815,182-65,815,318) and p21 promoter are negative controls. **(F,G)** Decrease in H3K27me3 levels correlates with increase in H3K36me3: (F) ChIP-qPCR with H3K27me3 antibody shows decreased level at *HOXD13*-C region (gene body) only in EPC1-PHF1 expressing K562 cell line (Bar graph in maroon color), validating our ChIP-seq results in (A). (G) ChIP-qPCR with H3K36me3 antibody shows increased level at *HOXD13*-C region (gene body) only in EPC1-PHF1 expressing K562 cell line (Bar graph in maroon color). MyoD promoter is used as a negative control. (Values are a ratio of %Input of H4ac and H2B) (n=2, error bars are range of the values). (n=2, error bars represent Range of the values). See also Figure S5 (B,C). **(H,I)** The H3K27me3 activity of EPC1-PHF1 is inhibited by the presence of H3K36me3. *In vitro* Histone Methyltransferase assay and Histone acetyltransferase assay on recombinant nucleosomes (rNCP) with or without H3K36me3. Purified complexes were normalized using western blotting for EZH2 and KAT5 (Fig S5E-F). Coomassie stained SDS-PAGE gel is the loading control for recombinant nucleosomes. **(J)** Schematic representation of the Topologically Associated Domains (TADs) and sub domains at the *HOXD* locus in mammalian cells. **(K)** Alignment of Hi-C data in K562 cells (Rao et al., 2014) with CTCF ChIP-seq in K562 cells (ENCODE/Broad histone marks) and ChIP-seq from this study (Figure 4A). Highlighted regions indicate histone modification changes confined to a topological domain in EPC1-PHF1 expressing cell line compared to tag-only expressing cell line (Mock). See also Figure S6.

Replacement of canonical H2A with H2A.Z in chromatin is another enzymatic activity of the NuA4/TIP60 complex through its EP400 subunit (Billon and Cote 2013; Pradhan et al. 2016). ChIP-qPCR with a H2A.Z specific antibody revealed a similar mislocalization of this variant histone at the *HOXD* sites (Figure 3E). These data clearly validate our hypothesis that the EPC1-PHF1 fusion complex can localize at genomic regions normally occupied by PRC2, mislocalizing the activities of the TIP60 complex to deregulate transcription. Other genes/genomic loci normally targeted by PRC2 and bound by the fusion show clear *de novo* H4 acetylation and decrease of H3K27me3 (Figure S5D).

We were intrigued at the apparent reduction in the levels of H3K27me3 at the *HOXD* gene cluster in the EPC1-PHF1 expressing cells (Figure 3A-highlighted in yellow). As shown above, we observed neither a global decrease in H3K27me3 levels (Figure S2E) nor destabilization of the PRC2 complex in our cell line (Figure 1). Moreover, we could detect productive histone methylation by the EPC1-PHF1 mega-complex (Figure 1H, Figure S2D). In a previous study (Musselman et al. 2012), we demonstrated that the Tudor domain of PHF1 binds to H3K36me3 and constrains PRC2 mediated H3K27me3 activity. We hypothesized that this could be part of the mechanism leading to a local decrease of H3K27me3. We checked for the levels of H3K27me3 and H3K36me3 at the *HOXD13*-C locus by ChIP-qPCR to confirm this possible cross-talk. A decrease in H3K27me3 was observed in EPC1-PHF1 cells compared to controls, validating our ChIP-seq results (Figure 3F). We also saw an increase in H3K36me3 at the *HOXD13*-C locus in EPC1-PHF1 cells (Figure 3G), likely linked to the increased transcription of the gene. This cross-talk was not observed at the NuA4/TIP60-bound and highly transcribed gene *RPSA* (Figure S5B)(Jacquet et al. 2016) or at *HOXA9*, a repressed gene not occupied by the EPC1-PHF1 fusion complex (Figure S5C). We conclude that the mislocalization of H4 acetylation and the variant histone H2A.Z at the posterior *HOXD* gene locus in cells expressing EPC1-PHF1 leads to de-repression and productive transcription. Subsequent deposition of H3K36me3 – a histone mark linked to transcription elongation - blocks the spread of the repressive H3K27me3 mark, possibly in part through its recognition by the Tudor domain of PHF1 and direct effect on EZH2/PRC2 activity (Schmitges et al. 2011; Musselman et al. 2012; Jani et al. 2019; Finogenova et al. 2020). We confirmed this cross-talk *in vitro* with our purified fractions in HMT and HAT assays using recombinant nucleosomes carrying the H3K36me3 mark (Figure 3H-I, S5E-F). Both PHF1/PRC2.1 and EPC1-PHF1 complexes are clearly inhibited in their methyltransferase activity by the presence of H3K36me3, while TIP60 and EPC1-PHF1 complexes are not affected in their acetyltransferase activity.

### The EPC1-PHF1 mediated effect on histone acetylation/methylation at *HOXD* is restricted to a specific Topologically Associating Domain (TAD)

The *HOXD* gene cluster is involved in mammalian axial patterning, limb and genital development. Spatiotemporal regulation of the *HOXD* locus depends on long-acting multiple enhancer sequences located in “gene deserts” outside the *HOXD* cluster. Mammalian cells organize such regulatory landscapes in structural units called Topologically Associating Domains (TADs), which maintain genomic contacts even in the absence of transcription. These pre-formed 3D structures, delimited by CTCF and cohesin proteins, remain globally similar in cell types and are conserved from mouse to humans (reviewed in (Bompadre and Andrey 2019; Szabo et al. 2019)). The *HOXD* locus is positioned between two large TADs (Rodriguez-Carballo et al. 2017)(Figure 5A). During limb development, the telomeric TAD is activated and controls the transcription of early *HOXD* genes (forearm), while later the centromeric TAD controls the posterior *HOXD* genes (digits)(Andrey et al. 2013). A strong boundary limits the posterior *HOX* genes from getting activated aberrantly (Lonfat and Duboule 2015).

Chromatin modifications influence the dynamic mammalian TADs. The repressive H3K27me3 modification segregates chromatin into discrete sub-domains, and studies suggest that these 3D clusters require PcG proteins and H3K27me3 (Szabo et al. 2019). The *HOXD* cluster is known to harbor an H3K27me3 dense region, and the 1.8kb region between *HOXD11* and *HOXD12* (D11.12) may act as a putative mammalian Polycomb Responsive Element (PRE) (Woo et al. 2010). Recent studies have shown that this region acts as a nucleation hub for PRC2 to stably bind and spread H3K27me3 through intra- and interchromosomal interactions (Vieux-Rochas et al. 2015; Oksuz et al. 2018).

We observed that the Tip60/KAT5-mediated H4 acetylation and reduction of H3K27me3 at the *HOXD* locus in EPC1-PHF1 cells localized to the posterior *HOXD* genes. This led us to explore the possibility that a TAD boundary exists in this region. When we aligned the publicly available CTCF ChIP-seq data and Hi-C dataset in K562, we observed that the chromatin changes induced by the EPC1-PHF1 fusion protein are restricted to a large portion of the centromeric TAD at the *HOXD* locus (Figure 3J-K). Since Polycomb group protein-mediated long-range contacts are also at play at the *HOXD* locus, we looked for spreading of EPC1-PHF1-mediated H4 acetylation to other genomic loci that were shown to interact with the *HOXD* locus. Although existing Hi-C/5-C studies in K562 cells are not detailed enough to detect PcG-mediated TADs (Kundu et al. 2017) and many of the explored regions are not conserved from mouse to humans (Vieux-Rochas et al. 2015; Oksuz et al. 2018), we could detect the presence of H4 acetylation at the *HOXC* locus (H3K27me3 spreading site), *EVX1*, *HOXB13*, *CYP26B1* and *LHX2* (H3K27me3 nucleation sites) (Figure S6). This observation suggests that the EPC1-PHF1 fusion protein could mislocalize activating histone marks through PcG-mediated chromatin contacts, activating an oncogenic transcriptional network. A similar mechanism was observed recently for an oncogenic EZH2 mutant that co-opted PcG-mediated long-range chromatin contacts to repress multiple tumor suppressors (Donaldson-Collier et al. 2019). Further work will be required to fully dissect the effect of the EPC1-PHF1 fusion complex on the structure and dynamics of TADs/polycomb-mediated chromatin contacts.

### JAZF1 is a stochiometric interaction partner of the NuA4/TIP60 complex

Various subunits of the NuA4/TIP60 complex can be fused to different subunits from Polycomb Repressive complexes 1 and 2 in Endometrial Stromal Sarcoma (Figure 1A). However, the most frequently observed translocation is that of *JAZF1-SUZ12,* which occurs in more than 50% of all low grade ESS (LG-ESS (Ferreira et al. 2018)). There are other recurrent translocations involving the JAZF1 protein in sarcomas, namely *JAZF1-PHF1* and *JAZF1-BCORL1* (Micci et al. 2006; Schoolmeester et al. 2013; Allen et al. 2017)(Figure 1A). This connection led us to investigate whether JAZF1 interacts with the TIP60 complex.

The molecular function of the JAZF1 protein is largely unexplored. It is a transcription factor with three C2H2 zinc fingers, and studies have linked it to glucose production, lipid metabolism, ribosome biogenesis, metabolic disorders and pancreatic cancer (Nakajima et al. 2004; Liao et al. 2019; Kobiita et al. 2020; Zhou et al. 2020). The JAZF1 protein has a homolog in *S. cerevisiae* called Sfp1 (Figure S7A), an essential transcription factor involved in regulation of ribosomal protein (RP), ribosomal biogenesis(RiBi) and cell cycle genes, responding to nutrients and stress through regulation by TORC1 (Marion et al. 2004; Lempiainen et al. 2009). Crucially, Sfp1 and the yeast NuA4 complex cooperate functionally to regulate RP and RiBi genes (Figure S7B) (Reid et al. 2000; Loewith and Hall 2011; Rossetto et al. 2014).

Since *JAZF1* is not expressed in K562 cells (www.proteinatlas.org), we performed multiple cell compartment affinity purification coupled to tandem mass spectrometry (MCC-AP-MS/MS) in HEK293 as described in (Lavallee-Adam et al. 2013). This technique allowed us to isolate protein-protein interaction occurring on the chromatin fraction. By purifying JAZF1, we recovered several NuA4/TIP60 core components with good reliability (Figure S7C). JAZF1 is highly expressed in the male and female reproductive tissues and may thus be involved in the tumorigenesis of LG-ESS of all fusion subtypes. To confirm this interaction, we ectopically expressed JAZF1-3xFLAG-2xSTREP in K562 cells from the *AAVS1* locus and tandem affinity-purified the protein from nuclear extracts. Silver staining of the purified fraction on gel, western blot and mass spectrometry analyses all confirm a very tight association of JAZF1 with the NuA4/TIP60 complex (Figure 4A-B, S7D), suggesting that JAZF1 can be a stable stoichiometric subunit. Accordingly, JAZF1 occupies the promoters of known TIP60-bound Ribosomal protein genes *RPSA* and *RPL36AL* as determined by ChIP-qPCR (Jacquet et al. 2016) (Figure 4D).

**Figure 4:**
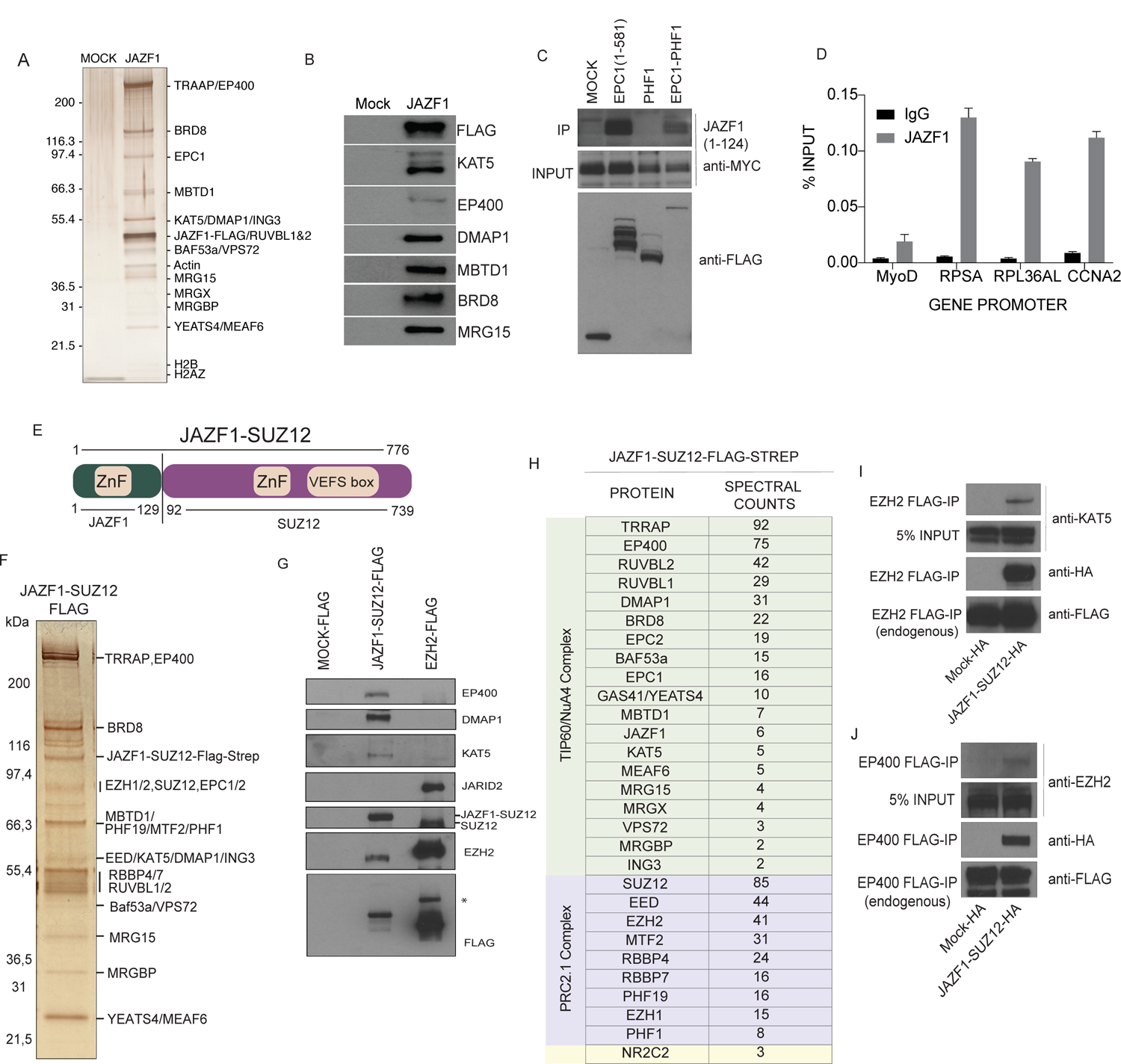
The JAZF1 protein stably associates with the NuA4/TIP60 complex. **(A)** Silver stained SDS-PAGE showing tandem affinity-purified JAZF1. Labels on the right side are proteins that were identified based on western blotting and predicted molecular weights. The JAZF1 fraction shows all subunits of the NuA4/TIP60 complex. See also Figure S7C-D for Mass Spectrometry analysis of JAZF1 purification from HEK293 and K562 cells, respectively. **(B)** Western blots of selected NuA4/TIP60 subunits on the affinity purified fraction shown in (A). **(C)** Immunoprecipitation of JAZF1 with the EPC1-PHF1 fusion. JAZF1 (1-124) corresponding to the portion in the JAZF1-SUZ12 fusion protein interacts with EPC1(1-581) and EPC1-PHF1. See Figure S7E for data with full length JAZF1 construct. **(D)** FLAG-JAZF1 ChIP-qPCR. Anti-FLAG ChIP-qPCR in HEK293 cells stably expressing FLAG tagged JAZF1. Enrichment of JAZF1 can be seen at ribosomal protein gene promoters (*RPSA* and *RPL36AL*) regulated by NuA4/TIP60. *CCNA2* is a positive control for JAZF1 (Values are represented as %IP/input chromatin, n=2, Error bars represent range of the values). **(E)** Schematic representation of the JAZF1-SUZ12 fusion protein. The numbers indicated are amino acids, protein domains retained in the fusion are indicated. **(F)** Silver stained SDS-PAGE showing tandem affinity-purified JAZF1-SUZ12. Labels on the right side are proteins that were identified based on western blotting and predicted molecular weights. **(G)** Western blots of selected NuA4/TIP60 and PRC2 subunits on the affinity purified fraction shown in (F). Note the absence of JARID2 in the JAZF1-SUZ12 complex. **(H)** Mass Spectrometry analysis of the affinity purified JAZF1-SUZ12 complex shown in (F). Note the absence of JARID2 and the presence of the 3 polycomb-like paralogs PHF1, MTF2 and PHF19, identifying PRC2.1. **(I,J)** Formation of a TIP60-PRC2 mega-complex by JAZF1-SUZ12. (I) Immunoprecipitation of endogenous tagged EZH2, Tip60 can be detected only in the JAZF1-SUZ12 expressing cells. (J) Immunoprecipitation of endogenous tagged EP400, EZH2 can be detected only in the JAZF1-SUZ12 expressing cells.

To analyze this tight interaction in the context of the *JAZF1-SUZ12* translocation, we performed co-immunoprecipitation experiments showing that the TIP60 complex still interacts strongly with the portion of JAZF1 (JAZF1(1-124)) that is fused to SUZ12 (Figure 4C, S7E). The same experiments showed that JAZF1 also still interacts with EPC1(1-581) and the EPC1-PHF1 mega-complex, but not with PHF1/PRC2.1. We then produce an isogenic cell line expressing the JAZF1-SUZ12 oncogenic fusion and purified the associated protein complex. Analysis of the purified fraction on gel, by western blot and mass spectrometry again shows the assembly of a chimeric mega-complex merging NuA4/TIP60 to PRC2.1 (Figure 4E-H), confirmed again by fractionation of endogenous respective complexes (Figure 4I-J). These conclusive data indicate that JAZF1 fusions could have the same effects on the epigenome as EPC1-PHF1.

### Conserved oncogenic mechanism of EPC1-PHF1 and JAZF1-SUZ12 fusion proteins

We have shown that the oncogenic EPC1-PHF1 fusion complex can lead to local transcription activation despite associating with a functional PRC2.1 complex. To further confirm the effect on transcription, we used a chemically inducible dCas9-based site-specific recruitment assay in HEK293T cells (Alerasool et al. 2022)(Figure 5A). In this system we could detect specific transcriptional activation by the EPC1 protein, EPC1(1-581) and EPC1-PHF1, but not by PHF1 (Figure 5B). Interestingly, we found that the JAZF1 protein is a very robust transcriptional activator in this reporter system. This is also the case for the JAZF1-SUZ12 fusion protein, as well as for the truncated JAZF1 (1-124), but not SUZ12 by itself (Figure 5C). ChIP-qPCR of H4 acetylation at the reporter gene also specific increase with the two fusions but not with PHF1 (Figure S8A-B).

**Figure 5:**
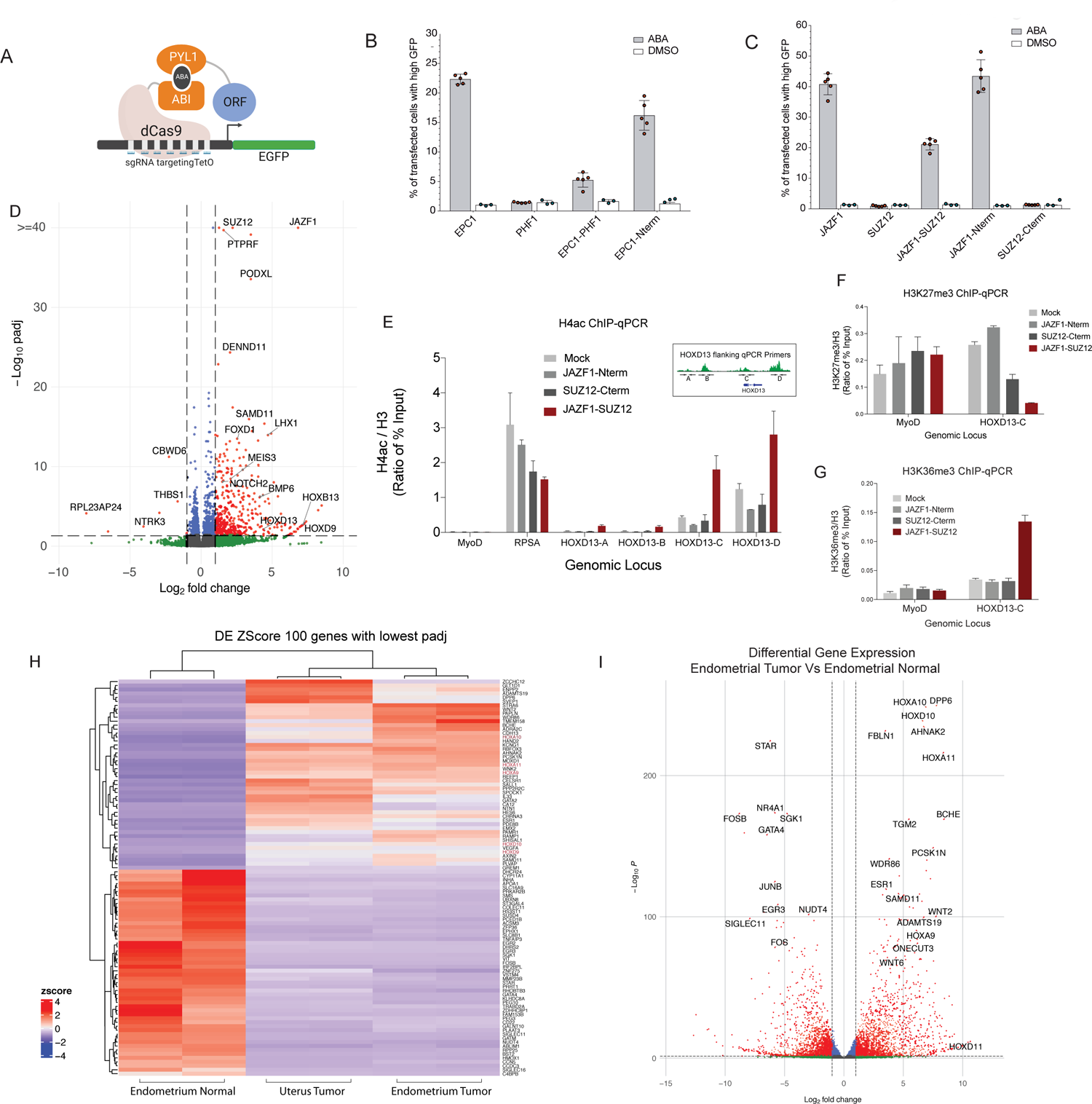
The JAZF1-SUZ12 fusion protein has molecular impacts like EPC1-PHF1. **(A)** Model for the dCas9-based inducible reporter gene activation assay in (B-C). **(B,C)** Transcription activation assay quantified by Flow cytometry analysis. Panel B shows the % of cells transfected with EPC1, PHF1, EPC1-PHF1 or EPC1 (1-581) expressing high GFP upon ABA treatment. EPC1-PHF1 is a transcriptional activator like EPC1 and EPC1-N terminus, whereas PHF1 is not. Panel C shows the % of cells expressing high GFP when transfected with JAZF1, SUZ12, JAZF1-SUZ12, JAZF1(1-129) and SUZ12(93-740) and treated with ABA. JAZF1 (Full length and N terminus) and JAZF1-SUZ12 act as transcriptional activators whereas SUZ12 does not. At least 25,000 cells were analyzed for each replicate. Error bars represent SEM of 5 independent repeats. DMSO is used as a negative control. **(D)** Volcano plot of differential gene expression analysis of K562 cell line expressing JAZF1-SUZ12 fusion compared to empty vector K562 cell line. **(E)** Mislocalization of H4 acetylation determined by ChIP-qPCR. JAZF1-SUZ12 expressing K562 cell line (Bar graph in maroon color) shows increased H4 acetylation compared to control cell lines at *HOXD13*-A, B, C, D regions, like EPC1-PHF1 in Figure 4C (Values are represented as ratio of % of input chromatin of H4ac and H3) (n=2, error bars represent Range of the values). *RPSA* promoter is a positive control for H4 acetylation, *MYOD* promoter is a negative control. **(F, G)** Decrease in H3K27me3 levels at the *HOXD* locus correlates with increased H3K36me3 in JAZF1-SUZ12 expressing cells: **(F)** ChIP-qPCR with H3K27me3 antibody shows decreased level at *HOXD13*-C region (gene body) in JAZF1-SUZ12 expressing K562 cell line (Bar graph in maroon color). *MyoD* promoter is a negative control. **(G)** ChIP-qPCR with H3K36me3 antibody shows increased level at *HOXD13*-C region (gene body) only in JAZF1-SUZ12 expressing K562 cell line (Bar graph in maroon color). *MyoD* promoter is a negative control. (Values are represented as ratio of % of input chromatin of H3K27me3 or H3K36me3 and H3) (n=2, error bars represent Range of the values). See also Figure S8D-E. **(H,I)** RNA sequencing followed by differential gene expression analysis of two Low Grade Endometrial Stromal Sarcoma patient tissue samples with presence of JAZF1-SUZ12 fusion. Pair-wise comparison was performed to an Endometrial normal patient tissue (paired normal of Endometrial Tumor sample). **(H)** Heatmap showing hierarchical clustering of top 100 differentially expressed genes. **(I)** Volcano plot of differential gene expression analysis of Endometrial Tumor with JAZF1-SUZ12 fusion compared to adjacent normal Endometrial tissue. See also Figure S10 and S11.

The gene expression profiles of ESS with different fusions (JAZF1 and TIP60) cluster together (Dewaele et al. 2014; Micci et al. 2016; Przybyl et al. 2018). We also performed an RNA-seq experiment with K562 cells expressing JAZF1-SUZ12 and found again specific increased expression of *HOX* genes/PRC2 targets, including *HOXD13* (Figure 5D, S9, Table S2). Recent interactome analyses of JAZF1-SUZ12 and JAZF1 also validate our results (Piunti et al., 2019)(Procida et al. 2021). Altogether, these data confirm that JAZF1 associates with the TIP60 complex and that all of the currently detected fusion proteins in LG-ESS bridge the TIP60 complex to Polycomb repressive complexes.

To confirm that the JAZF1-SUZ12 fusion generates the same molecular events on chromatin as EPC1-PHF1 we looked again at the *HOXD* locus by ChIP-qPCR. In JAZF1-SUZ12 expressing cells, we again detect an increase in H4 acetylation (Figure 5E, S8C), a decrease in H3K27 methylation (Figure 5F, S8D-E) and an increase in transcriptional elongation-associated H3K36me3 (Figure 5G, S8D-E) in the region flanking the *HOXD13* gene. JAZF1-SUZ12 is thus able to mislocalize the TIP60 complex-associated histone mark through the JAZF1 protein, drawing parallels with the mislocalization mediated by EPC1-PHF1 (Figure 3).

The loss of H3K27me3 in these data can be explained by the loss of the ZnB domain of SUZ12 in the JAZF1-SUZ12 fusion protein, which has been shown to reduce the binding of JARID2 and EPOP subunits (Chen et al. 2018a). SUZ12 acts as a platform to assemble the non-canonical subunits of the PRC2 complex, and a recent study shows that mutations in SUZ12 can modulate PRC2.1 vs PRC2.2 formation (Youmans et al. 2021). Furthermore, recent structural data suggest that the JARID2-containing PRC2.2 complex can partially override H3K36me3-mediated inhibition of EZH2 catalytic activity (Kasinath et al. 2021), whereas the PRC2.1 complex is known to be catalytically inhibited by the presence of H3K36me3 (Musselman et al. 2012; Finogenova et al. 2020). The loss of JARID2 interaction in JAZF1-SUZ12 skews it to associate with the PRC2.1 complex, as clearly shown by our purification (Figure 4G-H), which is catalytically inhibited by the presence of the H3K36me3 modification.

### Transcriptome analysis of human ESS tissue samples confirms upregulation of PRC2 target genes

In order to support the mechanistic model proposed by studying the fusions in our heterologous model system, we performed RNA-sequencing of two LG-ESS patient tissue samples with the *JAZF1-SUZ12* translocation and one paired normal endometrial tissue (Figure 5H-I, S10A, Table S3). Two pair-wise differential expression (DE) analysis were performed against the endometrial normal tissue sample. Pairwise differential expression analysis of endometrial tumor with the adjacent normal tissue is presented as a volcano plot in Figure 5I (the other ESS sample is shown in S10A). Significantly upregulated genes include many homeobox genes such as *HOXA10, HOXD10, HOXA11, HOXD11* and *HOXA9*, showing that fusion proteins in LG-ESS target the *HOX* gene cluster and drawing parallels to our result from K562 cell lines. Different pathway enrichment analyses clearly identify genes regulated by Polycomb Group proteins and H3K27me3 as upregulated in these endometrial tumors (Figure S10 and S11, Table S3), hinting that the JAZF1-SUZ12 fusion complex mis-targets activating marks at these genes. Altogether, our data strongly support a model that the mislocalization of TIP60 activities could be the predominant oncogenic mechanism of the recurrent fusion proteins found in Endometrial Stromal Sarcomas and also other Sarcomas.

## DISCUSSION

Our detailed biochemical work demonstrates that the EPC1-PHF1 and JAZF1-SUZ12 fusion proteins found in sarcoma assemble a mega-complex that merges two important chromatin modifying activities with opposite functions in gene expression. Our results show that binding of this EPC1-PHF1/JAZF1-SUZ12 chimeric complexes increase histone acetylation and H2A.Z incorporation within regions regulated by PRC2, notably within the *HOXD* gene cluster. This is correlated with increased gene expression, consistent with the notion that the fusions mistarget the HAT and histone exchange activities of NuA4/TIP60 to chromatin regions that are normally maintained in the silenced state (Figure 6). Interestingly, mistargeting of histone acetylation at the *HOXD* cluster leads to loss of H3K27 trimethylation over an entire specific topology-associating domain (TAD) without affecting the neighboring ones, albeit also bound by PRC2 complexes. In theory, since the fusions are integrated in a mega-complex with PRC2.1, it could act at most PRC2.1 binding sites. We speculate that the presence of small preexisting TIP60 binding sites within or in the vicinity of Polycomb/repressed regions, as we see at the *HOXD* cluster, could make a big difference. The TIP60 moiety of the EPC1-PHF1 mega-complex could be efficiently recruited there (by DNA-bound factors and/or small regions of H3K4me3 in bivalent chromatin) while the PRC2 moiety would allow it to spread along the region like PRC2 normally does, leading to de novo acetylation. This would lead to increased transcription, coupled with H3K36 tri-methylation, which in turn inhibits the HMT activity of PRC2 (through recognition by PHF1 Tudor domain), leading to a decrease of H3K27me3 over the region (Figure 6, middle panel). It is important to note that H3K36 methylation has also been described as a potent boundary mark to block H3K27me3 spreading (Streubel et al. 2018). This mechanism would also explain why the clearer changes in histone marks and gene expression are found at EPC1-PHF1 mapped binding sites that overlap with both TIP60 and PRC2.1 (Figure 2).

**Figure 6:**
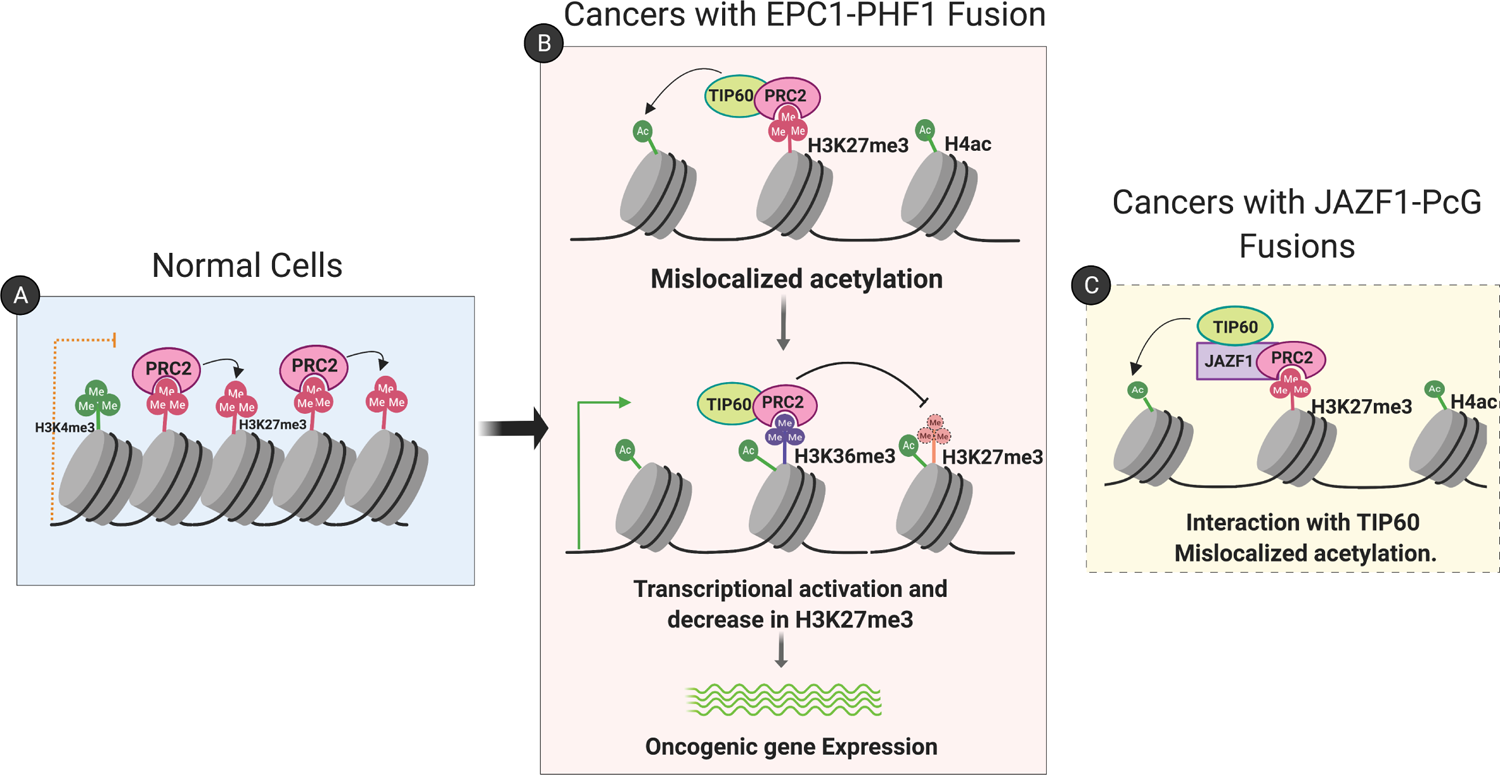
Model for the oncogenic mechanism of EPC1-PHF1 fusion protein in soft tissue sarcomas. **(A)** In normal cells, PRC2 complex occupies repressed or poised chromatin. **(B)** When EPC1-PHF1 fusion protein is expressed, it assembles a mega-complex combining NuA4/TIP60 and PRC2 complexes. The mega-complex occupies regions that have activating and repressive histone marks (such as the *HOX* clusters in K562) and mislocalizes NuA4/TIP60 activities (H4, H2A acetylation, H2A.Z exchange). This tips the balance towards transcriptional activation. Levels of the transcription elongation associated histone mark H3K36me3 increase and inhibits the deposition of the repressive H3K27me3 histone mark. Thus, the changes in chromatin landscape potentiate the expression of oncogenes. Sarcomas with fusions of JAZF1 and PcG proteins also employ a similar mechanism since JAZF1 strongly interact with the NuA4/TIP60 complex and mislocalize its activities.

In addition, our study validates the protein JAZF1 as a potent transcription activator that stably associates with the NuA4/TIP60 complex. This information further integrates all of the reported translocations in Low-grade Endometrial Stromal sarcomas (LG-ESS) as producing a physical merge of TIP60 and PcG complexes, unifying the underlying molecular mechanism (Figure 6, right panel). This is supported by the robust clustering of gene expression profiles of the LG-ESS with various fusion proteins when compared to High-grade ESS or tumors without fusion genes (Micci et al. 2016). Our findings further underline the importance of deregulated epigenetic modifiers in promoting oncogenesis, particularly in sarcomas (Nacev et al. 2020), in part through changes of the chromatin landscape over large genomic regions. The highly recurrent physical merger of TIP60 and PcG complexes in these sarcomas, through several distinct subunits, indicates that this event is the driving force in the oncogenic mechanism. It is somewhat reminiscent of the MLL fusions proteins in pediatric acute myeloid leukemia, in which the H3K4 HMT MLL is recurrently fused with diverse components of the super-elongation complex, creating a large complex driving oncogenesis through altered transcription programs involving chromatin modifiers and readers as well as *HOXA* cluster misregulation (Mohan et al. 2010). A parallel can also be drawn with the discovery of various BRD4-linked Z4 complex proteins that are fused to NUT in NUT midline carcinoma (Shiota et al. 2018).

At the *HOXD* locus, the chromatin modification changes observed with EPC1-PHF1 spread over a large region that encompasses 2 domains within a larger TAD (Figure 3J-K). This is especially interesting given recent studies demonstrating the nucleation and spreading of H3K27me3 through Polycomb domain contacts (Vieux-Rochas et al. 2015; Oksuz et al. 2018). Interestingly, EPC1-PHF1 localizes to the *HOXD13*/*EVX2* nucleation region and mislocalizes H4 acetylation only in the centromeric TAD. We could also observe EPC1-PHF1 localization and H4 acetylation presence at “spreading” sites (Figure S5). A similar observation was made for the Ewing sarcoma fusion protein at the *HOXD* centromeric TAD leading to overexpression of *HOXD13* as seen here (Svoboda et al. 2014; von Heyking et al. 2016). A question that emerges from our observation is whether mislocalization of activating marks will lead to disruption of these H3K27me3-centric repressive domains and disrupt the organization of chromatin domains. Disruption of polycomb-mediated repression by fusion proteins to activate stem cell-related gene expression programs has emerged as a common theme recently (Brien et al. 2019). Unlike in Synovial sarcoma or NUT midline carcinoma, we do not observe extensive mislocalization/disruption in chromatin. This could be an artifact of our model system of study. Alternatively, the LG-ESS pathology is rare, milder, not rapidly progressing and with limited metastasis. LG-ESS may hence only deregulate few genes. Direct chromatin profiling in patient samples with the JAZF1-SUZ12 fusion will provide further evidence for this hypothesis.

Intriguingly, a newly described subunit of PRC2 complexes, EZHIP (CXORF67), was also found fused to the TIP60 subunit MBTD1 in LG-ESS (Dewaele et al. 2014). EZHIP functions similarly to the oncogenic H3K27M mutant by binding to EZH2 and inhibiting its activity in trans (Hubner et al. 2019; Jain et al. 2019; Piunti et al. 2019). This mechanism seems similar to the trans-inhibition of EZH2 through PHF1 Tudor-H3K36me3 binding and recently described direct inhibitory effect of H3K36me3 on EZH2 catalytic activity (Jani et al. 2019; Finogenova et al. 2020). The molecular characterization of MBTD1-EZHIP will further reveal if parallels can be drawn with EPC1-PHF1/JAZF1-SUZ12.

Since Endometrial Stromal sarcomas and OFMTs are very rare, we were fortunate to get access to patient tissues. Although limited by sampling size, this is the first report of a gene expression comparison of a LG-ESS sample to adjacent normal endometrial tissue. While the results clearly support the conclusions drawn from the model system, further analysis of more paired LG-ESS samples may shed light on the oncogenic signature of this rare cancer. Many pathways identified as enriched in our expression analysis of the two patient samples are implicated in oncogenesis of sarcomas (Figure S10, S11, Table S3)(Damerell et al. 2021).

The data presented in this study indicate that the mislocalization of NuA4/TIP60 leads to histone H4 acetylation and H2A.Z incorporation in chromatin, favoring local gene expression that can subsequently inhibits PRC2 activity. This succession of molecular events is likely the predominant oncogenic mechanism of the recurrent fusion proteins found in Endometrial Stromal Sarcomas, Ossifying Fibromyxoid Tumors, and other related Sarcomas (Figure 1A). Since the mutational burden of translocation carrying cancers is relatively lesser than other cancers (clonal homogeneity), a true characteristic of sarcomas, targeted therapeutic approaches specific to the fusion protein are plausible (Taylor et al. 2011; Brien et al. 2019). According to our study, targeting TIP60 HAT activity with available small molecule inhibitors seems a viable approach (Brown et al. 2016). Alternatively, development of inhibitors designed to disrupt the interaction between JAZF1 and the TIP60 complex may also be a promising therapeutic approach.

## MATERIALS AND METHODS

### Cell culture

K562 cells were obtained from the ATCC and maintained at 37°C under 5% CO2 in RPMI medium supplemented with 10% fetal bovine serum and GlutaMAX. 25 mM HEPES-NaOH (pH 7.4) was added during culture in Spinner flasks. HEK293 cells were obtained from the ATCC and maintained at 37°C under 5% CO2 in DMEM medium supplemented with 10% fetal bovine serum.

### Construction of Recombinant DNA

RT-PCR product from Total RNA isolated from surgically removed endometrial stromal sarcoma tumor was a kind gift from Dr. Fransesca Micci (Micci et al. 2006). The cDNA containing the entire truncation of EPC1 and a fused portion of PHF1 up to base pair (bp) 139, the included portion of PHF1 contained a BglI restriction site, which was then utilized to reconstruct the full length fusion gene using a clone of PHF1 obtained from Open Biosystems Products (Hunstville, AL). This construct was then subcloned into the different expression plasmids. Human cDNA of *JAZF1* was purchased from GE Healthcare. Open reading frame (ORF) of *JAZF1* was subcloned into vectors and used to construct the truncation and the JAZF1-SUZ12 fusion by PCR. All constructs were verified by sequencing, and their expression was tested by transient transfection.

### Cell Line Generation

EPC1-PHF1 and controls were targeted to the *AAVS1* safe harbour locus in K562 cells, using Zinc Finger Nuclease (Hockemeyer et al. 2009) as described previously in (Dalvai et al. 2015).

### Purification of Complexes

Native complexes were purified as described in detail before in (Dalvai et al. 2015; Doyon and Cote 2016). The purified complexes were loaded on NuPAGE 4–12% Bis-Tris gels (Invitrogen) and visualized by silver staining. Fractions were then analyzed by mass spectrometry (details in supplementary methods).

### Affinity Purification followed by Immunoblotting

K562 cell lines were expanded to get 6-8 million cells. Cells were collected and washed twice in 1X PBS (Phosphate Buffered Saline), lysed for 30 min in 2X volume Lysis Buffer (450mM NaCl, 10% Glycerol, 50mM Tris-HCl pH8, 1%Triton-X 100, 2mM MgCl2, 0.1mMZnCl2, 2mM EDTA, 1mM DTT, Protease inhibitors), same volume of lysis buffer without salt was added to get a final salt concentration of 225mM. Fractions were centrifuged to prepare whole cell extracts. The extracts were incubated with FlagM2 agarose resin (Sigma) for 4hr at 4°C. The resin was centrifuged, washed in Lysis buffer with 225mM salt and Eluted with 3X Flag peptide (Sigma). The eluted fraction was loaded on 4-15% gradient gels with Input and immunoblotted with appropriate antibodies.

### ChIP and ChIP-sequencing

FLAG ChIPs were performed according to the protocol described in (Jacquet et al. 2016). Histone modification ChIPs were performed according to the protocol described in (Lalonde et al. 2013). Libraries for sequencing were prepared as described (Jacquet et al. 2016). Samples were sequenced by 50 bp single reads on HiSeq 2000 platform (Illumina). Reads from ChIP-seq experiments were obtained from the Genome Innovation Center at McGill University. Raw reads were trimmed using fastp v0.20.1 (Chen et al. 2018b). Quality check was performed on raw and trimmed data to ensure the quality of the reads using FastQC v0.11.7 (Andrews 2010) and MultiQC v1.5 (Ewels et al. 2016). Trimmed reads were aligned on the human genome (hg18) using bwa mem v0.7.17 (Li 2013) and samtools v1.13 (Li et al. 2009). Raw signal tracks and normalized tracks (in reads per million, RPM) were produced from mapped reads using deepTool’s v2.17.0 bamCoverage tool (Ramirez et al. 2014) and bedtools genomecov tool (Quinlan and Hall 2010), respectively. Tracks were converted to the bigwig format using bedGraphToBigWig v2.8 (Kent et al. 2010). The macs2 v2.2.1 software (Feng et al. 2012) was used to perform the peak calling. Region annotation and integration of ChIP-Seq data with Microarray data were performed using the ChIPseeker package v1.26.2 (Yu et al. 2015) in R v4.0.3 (Team 2013). The metaplots and graphical representations were produced with the ggplot2 v3.3.5 package (Wickham 2011). Boxplot statistics were computed using bins; n= number of regions analyzed where 1 region is 100 bins. The heatmaps were generated using ComplexHeatmap v2.6.2 (Gu et al. 2016) and EnrichedHeatmap v1.20.0 packages (Gu et al. 2018). The CUT&RUN files used in the heatmap were lift over to UCSC hg18, using UCSC lift over tool (Kuhn et al. 2013). Quantitative real-time PCRs were performed on a LightCycler 480 (Roche) with SYBR Green I (Roche) to confirm the specific enrichment at defined loci. The error bars represent standard errors based on two independent experiments. Quantitative real-time PCR primers used are listed in Supplementary Table 1.

### CUT and RUN sequencing

The experiment was performed as detailed in (Skene et al. 2018) with some modifications. 0.5 million K562 cells were pelleted, resuspended in 1XPBS. Cells were crosslinked with 0.1% formaldehyde for 1 min. Crosslinked cells were washed at room temperature and bound to 10 μl of Concanavalin A beads slurry. Cell and bead suspension was incubated in buffer containing 0.05% digitonin and 0.5ug of anti-HA (Epicypher 13-2010) antibody overnight at 4°C. Afterward, the beads were washed with digitonin buffer resuspended in buffer containing pAG-MNase (CUTANA EpiCypher, 1:20 dilution) and digitonin, incubated 10 min with agitation. Beads were washed and resuspended in ice-cold digitonin buffer. 1mM CaCl₂ was added and incubated for 2hrs at 4°C. The reaction was stopped by addition of stop buffer containing EDTA and EGTA, incubated at 37°C for 10 min. Supernatant containing released chromatin was separated and decrosslinked at 55°C overnight. DNA was purified with NEB Monarch PCR & DNA purification kit following the protocol enriching for short DNA fragments. DNA was quantified by Qubit HS DNA kit. Library was prepared following the manufacturer’s instruction in NEBNext Ultra II DNA kit for low input ChIP. 100bp paired end sequencing was performed with the Illumina NovaSeq 6000 system. Raw reads were trimmed using fastp v0.21.0 (Chen et al. 2018b). Trimmed reads were aligned on the human genome (hg38) using bwa mem v0.7.17 (Li 2013) and samtools v1.13 (Li et al. 2009). Raw signal tracks and normalized tracks (in reads per million, RPM) were produced from mapped reads using deepTool’s v2.17.0 bamCoverage tool (Ramirez et al. 2014) and bedtools genomecov tool (Quinlan and Hall 2010), respectively. Tracks were converted to the bigwig format using bedGraphToBigWig v2.8 (Kent et al. 2010). The macs2 v2.2.1 software (Feng et al. 2012) was used to perform the peak calling, with the following parameters: no lambda, fragment size to 14, mfold from 5 to 50, and keep of all duplicated tags.

### HAT and HMT Assays

Fractions of purified complexes were assayed for enzymatic activity on short oligonucleosomes and free histones isolated from HeLa S3 cells as described previously (Utley et al. 1996; Doyon et al. 2004; Musselman et al. 2012).

#### Histone Acetyltransferase (HAT) Assay

*In vitro* histone acetylation assay 0.5µg of the indicated substrates in a 15µl reaction containing 50mM Tris-HCl pH8.0, 10% glycerol, 1mM EDTA, 1mM DTT, 1mM PMSF, 10mM sodium butyrate and 0.25 µCi/µl (4.9 Ci/mmol) of [3H]-labeled Acetyl-CoA (Perkin Elmer) or 0.15mM unlabeled Acetyl-CoA (Sigma). Samples were either spotted on P81 membrane (GE Healthcare) for count or analyzed on 18% SDS-PAGE. For radioactive gel assays, Coomassie staining was followed by treatment with Enhance® (Perkin Elmer) and fluorography, or by western blot analysis for non-radioactive assays. In results from liquid counting, error bars represent standard deviation of independent reactions.

#### Histone Methyltransferase (HMT) Assay

*In vitro* histone acetylation assay 0.5µg of the indicated substrates in a 15µl reaction containing 20mM Tris-HCl pH8.0, 5% glycerol, 0.1mM EDTA, 1mM DTT, 1mM PMSF, 100mM KCl and 1 mCi of [3H]AdoMet (S-adenosyl-L-[methyl-3H] methionine, 80 Ci/mmol; SAM) (Perkin Elmer) or 0.15mM unlabeled S-adenosyl methionine for 45 min at 30°C. Samples were either spotted on P81 membrane (GE Healthcare) for Scintillation count or analyzed on 15% SDS-PAGE. For radioactive gel assays, Coomassie staining was followed by treatment with Enhance® (Perkin Elmer) and fluorography, or by western blot analysis for non-radioactive assays. In results from liquid counting, error bars represent standard deviation of independent reactions.

The histones acetyltransferase (HAT) assays and histones methyltransferase (HMT) assays with recombinant Nucleosome Core Particles (rNCP) were performed in a volume of 15ul using 0.5ug of reconstituted/recombinant H3.3 mononucleosomes (unmodified or H3.3K36me3, epicypher #16-0390). The amount of purified enzyme complex used was normalized by western blotting to obtain similar amount of TIP60 complex in HAT reactions or PRC2 complex in HMT reactions. (For HAT: 2ul of Mock, 2ul of EPC1-PHF1, 0.635ul of EPC1(1-581); For HMT: 5ul of Mock, 5ul of EPC1-PHF1,1.875ul of PHF1)

### Recruitment-activator assay

The assay was performed as described in (Alerasool et al. 2022), Briefly, HEK293T TRE3G-EGFP reporter cell line with ABI-dCas9 and a gRNA targeting seven tetO repeats in the TRE3G promoter was generated and a clone showing robust EGFP induction by a strong transcriptional activator VPR was selected for downstream assays. 96-well plates were seeded with 3×10^4^ cells per well one day prior to transfection. 150 ng of each construct was transfected using polyethylemine (PEI). Transfected cells were induced 24 hours after transfection by treatment with 100 µM abscisic acid. 48 hours after induction, cells were dissociated and resuspended in flow buffer using a liquid handing robot and analyzed by LSRFortessa (BD). Flow cytometry data was analyzed using FlowJo by gating for positive gRNA (EBFP2), then further for construct (TagRFP) expression. At least 25,000 cells were analyzed for each replicate.

### Microarray

RNA samples were extracted using TRIzol reagent (Invitrogen), following manufacturer’s instructions. Duplicate RNA samples from EPC1-PHF1 expressing and control K562 cell lines were compared. We performed gene expression microarray experiments using the Human Illumina HumanHT-12_V4 platform. We first log2 transformed and quantile normalized the data before using them for the analyses presented in the paper. All the data were pre-processed using the lumi Bioconductor package (Du et al. 2008).

### Patient Tissue Samples

This study was approved by the Research Ethics Board of the University Health Network in Toronto, ON, Canada. Low Grade Endometrial Stromal Sarcoma biobanked specimens (Frozen) were obtained with broad consent. Fluorescence in situ hybridization (FISH) was performed to identify rearrangement involving *JAZF1* and *SUZ12*.

### mRNA Sequencing

RNA from K562 cell lines was extracted using the Monarch Total RNA purification kit (NEB) according to manufacturer’s instructions. RNA from frozen tissue samples were extracted using the miRNEASY micro kit (QIAGEN). Only samples with RIN above 7 was used for library preparation using the NEBNext Ultra II Directional RNA library + NEBNext Poly(A) mRNA-dual index kit. Sequencing run was performed in Illumina NovaSeq 6000 system. 100bp Paired-end reads were trimmed using fastp v0.20.1 (Chen et al. 2018b). Quality check was performed on raw and trimmed data to ensure the quality of the reads using FastQC v0.11.7 (Andrews 2010) and MultiQC v1.5 (Ewels et al. 2016). The quantification was performed with Kallisto v0.46.2 (Bray et al. 2016) against the human genome (hg38). The volcano graphical representations were produced with Bioconductor package Enhanced Volcano (Blighe et al. 2018). Differential expression analysis was also performed using the DESeq2 v1.30.1 package (Love et al. 2014). All R analysis were done in R v4.0.3 (Team 2013). Fusion transcripts were identified using the STAR-fusion pipeline (Haas et al. 2019). GSEA and Enrichment was carried out using clusterProfiler version v4.2.2 (Padj cutoff value = 0.05).

### Hi-C data alignment

Alignment of the ChIP sequencing data with previously published Hi-C data in K562 cells (Rao et al. 2014) was performed by converting the ChIP sequencing alignment from hg18 to hg19 using CrossMap Tool (Zhao et al. 2014). Hi-C data and ChIP sequencing data was then visualized using PyGenome Tracks tool V2.1 (Ramirez et al. 2018). The CTCF ChIP-sequencing data used is from GEO: GSM733719 (Broad Institute /ENCODE group). The Hi-C data (Heat map, domains, and loops) was downloaded from the datasets provided at Chorogenome (http://chorogenome.ie-freiburg.mpg.de/data_sources.html)

## Data And Software Availability

NGS assays reported in this study are available at the GEO repository under accession numbers below; ChIP-seq data and microarray expression analysis (<u>GSE162544),</u> CUT and RUN data (GSE196754), JAZF1-SUZ12 K562 RNA-seq (GSE196755), Patient sample RNA-seq (GSE196757) All Mass Spectrometry files generated as part of this study were deposited at MassIVE (http://massive.ucsd.edu). The MassIVE ID is MSV000083618, MSV000086476 and MSV000088749. The MassIVE FTP download links are ftp://massive.ucsd.edu/MSV000083618, ftp://massive.ucsd.edu/MSV000086476 and ftp://MSV000088749@massive.ucsd.edu. The password for download prior to final acceptance is “fusion”.

## COMPETING INTEREST STATEMENT

The authors declare no conflict of interest.

## ACKNOWLEDGMENTS

We thank Céline Roques, Valérie Côté and Philippe Cloutier for important technical support. We are very grateful to Prof. Francesca Micci for providing the cDNA obtained from patient sample that covered the EPC1-PHF1 fusion. We thank Compute Canada for the use of supercomputers, McGill Genome Center for sequencing/expression microarray and the CHUL proteomic platform. This work was supported by grants from the following: the Canadian Institutes of Health Research (CIHR) to J.C. (FDN-143314); the Government of Québec, Ministry of Economy and Innovation to B.C.; the Natural Sciences and Engineering Research Council (NSERC) of Canada to Y.D. (RGPIN-2014-059680) and J.-P.L. (RGPIN-2017-06124); University of Toronto startup funds to M.T. D.S., M.-E.L. and K.J. were supported by PhD studentships from Fonds de la Recherche Québec-Santé (FRQS) and Nature/Technologie (FRQNT). N.A. was supported by a CIHR post-doctoral fellowship and A.M. by MSc studentships from NSERC and FRQNT. B.C. holds the IRCM Bell-Bombardier Research Chair. A.-C.G. holds the Canada Research Chair in Functional Proteomics and the Lea Reichmann Chair in Cancer Proteomics. J.-P.L. and Y.D. are Junior 1 and Junior 2 FRQS scholars, respectively. J.C. holds the Canada Research Chair in Chromatin Biology and Molecular Epigenetics.

## AUTHOR CONTRIBUTIONS

D.S., N.A., M.-E.L., B.C., M.T., Y.D. and J.C. designed the experiments. D.S., N.A., M.-E.L., N.A., A.M., K.J., C.L., J.-P.L., J.R., J.L., and Y.D. performed the experiments. C.J.-B., E.P., L.H. and S.T.S. analyzed the genomic data. M.Q.B. and M.R. provided the patient samples. J.-P.L., A.-C.G., B.C., M.T., Y.D. and J.C. supervised and secured funding. D.S. and J.C. wrote the manuscript with the help of co-authors.

## Notes

### Competing Interest Statement

The authors have declared no competing interest.

