## Supplemental figures and methods for "Recurrent chromosomal translocations in sarcomas create a mega-complex that mislocalizes NuA4/TIP60 to Polycomb target loci"

Sudarshan et al.

**-SUPPLEMENTARY FIGURES AND LEGENDS**

**-SUPPLEMENTARY METHODS**

**-LIST AND CONTENT OF SUPPLEMENTARY TABLES (excel files)**

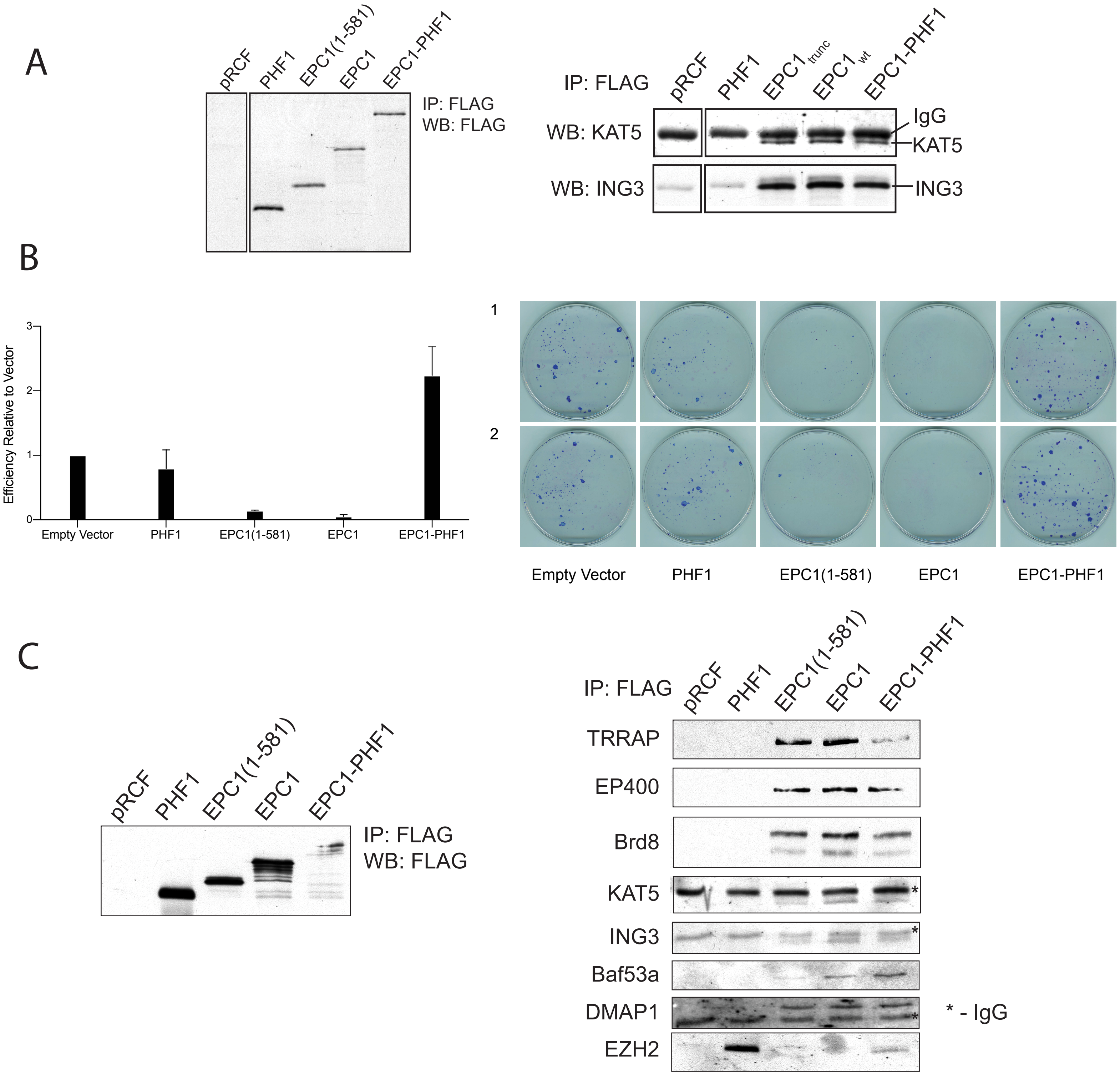
A
pRCF
PHF1
EPC1(1-581)
EPC1
EPC1-PHF1
IP: FLAG
WB: FLAG
IP: FLAG
pRCF
PHF1
EPC1trunc
EPC1wt
EPC1-PHF1
WB: KAT5
IgG
KAT5
WB: ING3
ING3
B
Efficiency Relative to Vector
0
1
2
3
Empty Vector
PHF1
EPC1(1-581)
EPC1
EPC1-PHF1
1
2
Empty Vector
PHF1
EPC1(1-581)
EPC1
EPC1-PHF1
C
pRCF
PHF1
EPC1(1-581)
EPC1
EPC1-PHF1
IP: FLAG
WB: FLAG
IP: FLAG
pRCF
PHF1
EPC1(1-581)
EPC1
EPC1-PHF1
TRRAP
EP400
Brd8
KAT5
ING3
Baf53a
DMAP1
EZH2
* - IgG

<u>Figure S1</u> **(Related to Figure 1). EPC1-PHF1 fusion protein associates with TIP60 as well as PRC2 complexes and enhances cell proliferation.**

**(A)** Indicated FLAG tagged constructs were transiently transfected in HEK293T and affinity purified by FLAG-resin after co-transfection with HA tagged KAT5 and ING3 and analyzed by western blotting. The left panel shows the normalized expression of transiently transfected FLAG tagged constructs. The right panel is the FLAG purification showing interaction with KAT5 and ING3.

**(B)** Colony formation assay in HEK293T cells. EPC1-PHF1 transfected cells show higher colony counts compared to controls. Upper panel shows quantification (n=2, error bars represent range of the values).

**(C)** FLAG affinity purification from stably transfected HEK293T cell lines (over-expression). The left panel shows relative expression levels of the indicated cell lines. The right panel is the western blot analysis of FLAG affinity purification showing interaction of EPC1-PHF1 fusion protein with endogenous TIP60 complex subunits (TRRAP, EP400, BRD8, KAT5, ING3, Baf53a, DMAP1) and EZH2.

A

| | Protein | Spectral Counts |
|---|---|---|
| TIP60 | TRRAP | 272 |
| | EP400 | 221 |
| | BRD8 | 61 |
| | EPC1 | 60 |
| | KAT5 | 52 |
| | DMAP1 | 60 |
| | ING3 | 31 |
| | RUVBL2 | 107 |
| | RUVBL1 | 98 |
| | BAF53a | 47 |
| | VPS72 | 23 |
| | MRG15 | 28 |
| | MRGX | 21 |
| | MRGBP | 12 |
| | GAS41/YEATS4 | 20 |
| | MEAF6 | 12 |
| | H2AZ1 | 5 |
| | H2BC3 | 5 |
| PRC2.1 | SUZ12 | 58 |
| | EZH2 | 39 |
| | EZH1 | 15 |
| | PHF1 | 57 |
| | EED | 25 |
| | RBBP4 | 18 |
| | RBBP7 | 15 |
| | PALI1 | 4 |

B

| | Protein | Spectral Counts |
|---|---|---|
| TIP60 | TRRAP | 57 |
| | EP400 | 28 |
| | BRD8 | 8 |
| | EPC1 | 9 |
| | KAT5 | 5 |
| | DMAP1 | 9 |
| | ING3 | 5 |
| | RUVBL2 | 25 |
| | RUVBL1 | 38 |
| | BAF53a | 8 |
| | ACTB | 7 |
| | VPS72 | 4 |
| | MRG15 | 3 |
| | GAS41/ YEATS4 | 4 |
| PRC2.1 | EZH2 | 3 |
| | PHF1 | 4 |
| | RBBP4 | 3 |

C

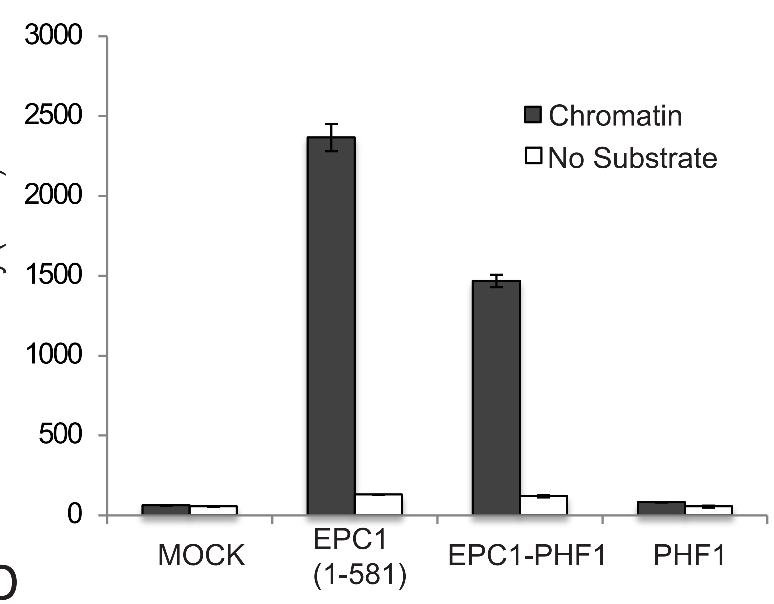


D

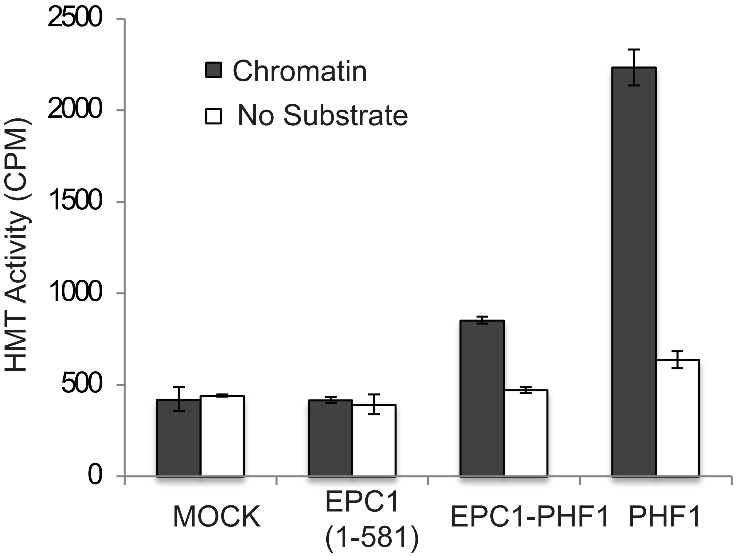


E

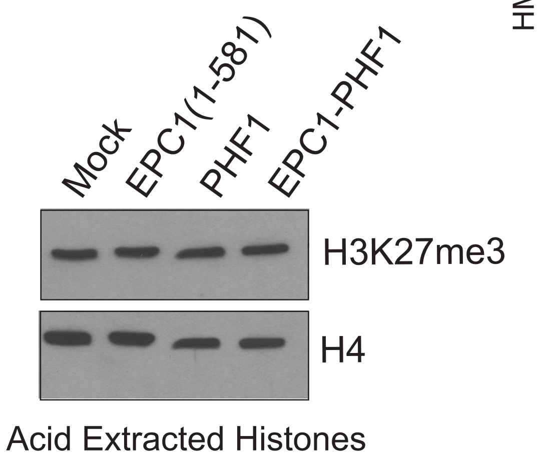

**<u>Figure S2</u> (Related to Figure 1). HAT and HMT activity of the mega-complex and the effect of EPC1-PHF1 on global H3K27me3 levels.**

**(A,B)** Mass Spectrometry analysis of the tandem affinity purified EPC1-PHF1 complex. Table with total spectral counts of peptides enriched in the affinity purification. Peptides highlighted in light-grey are NuA4/TIP60 complex components, peptides highlighted in darker-grey are PRC2.1 complex subunits. Replicates of Figure 1C-E using the same construct (A) or a EPC1-PHF1-3FLAG-HA-2A-puro cell line (B).

**(C)** Liquid HAT assay: Quantification of the Histone Acetyltransferase (HAT) assay in Figure 1H measuring the incorporation of $^3$H-Acetyl moiety on chromatin substrates by scintillation counting (Counts Per Minute/CPM, error bars are based on technical replicates). Reaction with purified EPC1-PHF1 complex (Figure 1C) shows elevated CPM, indicating HAT activity. EPC1 (1-581) is the positive control.

**(D)** Liquid HMT assay: Quantification of the Histone Methyltransferase (HMT) assay in Figure 2F measuring incorporation of $^3$H-Methyl moeity on chromatin substrates by scintillation counting (Counts Per Minute/CPM, error bars are based on technical replicates). Reaction with purified EPC1-PHF1 complex (Figure 1C) shows elevated CPM, indicating HMT activity. PHF1 fraction (PRC2) is the positive control.

**(E)** Immuno-blotting of acid extracted histones from EPC1-PHF1 and control cell lines (K562) with H3K27me3 antibody. H4 antibody is used as loading control for histones. Global level of H3K27me3 does not change in K562 with the expression of the EPC1-PHF1 fusion protein.

Sudarshan_Supplemental_FigS3

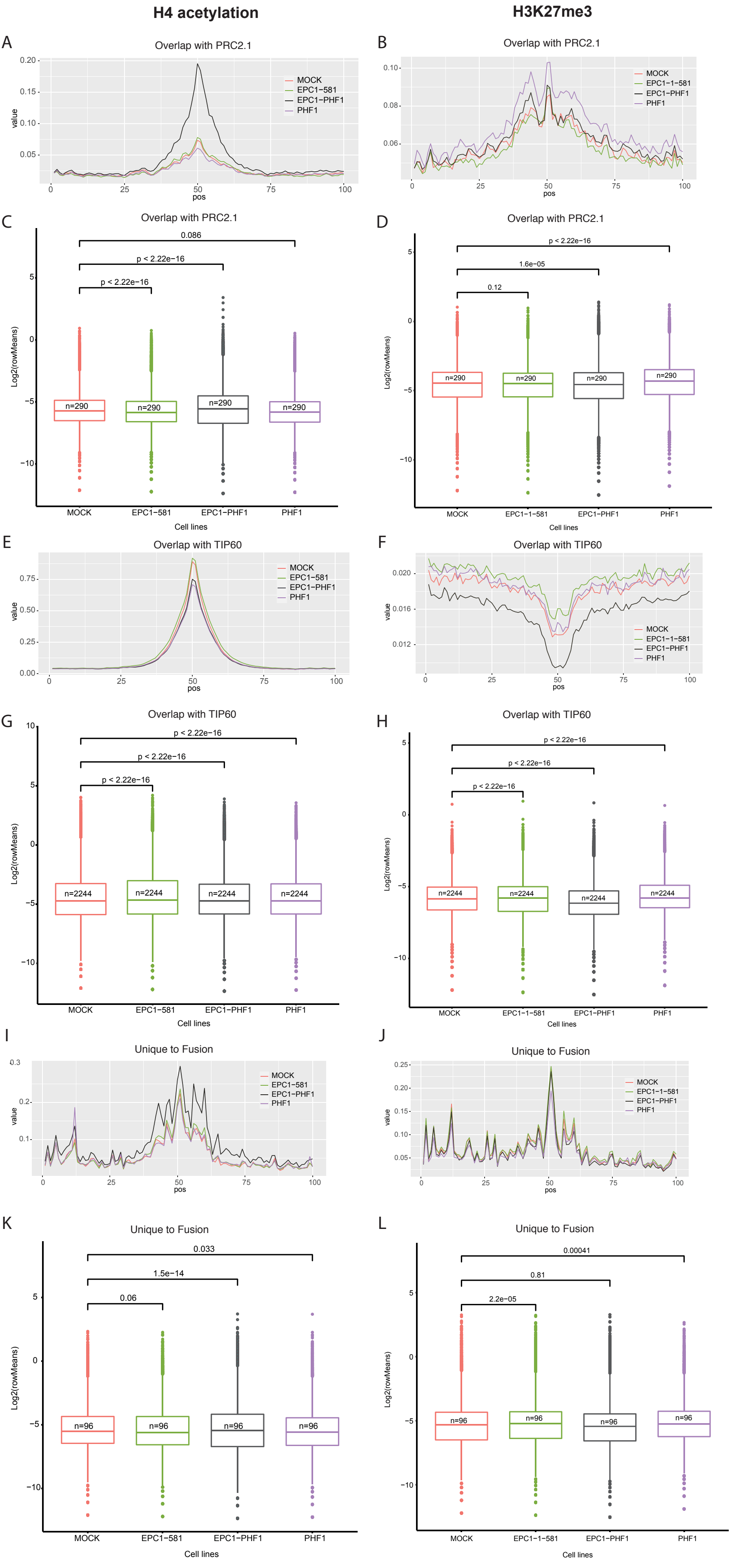

**<u>Figure S3</u> (Related to Figure 2): Histone marks changes at EPC1-PHF1 bound loci.**

**(A,B,E,F,I,J)** Density Plots of H4 acetyl and H3K27me3 enrichment in K562 cell lines in indicated partitions of EPC1-PHF1 peaks.

**(C,D,G,H,K,L)** Boxplots showing H4 acetyl and H3K27me3 enrichment level in K562 cell lines in indicated partitions. Statistics were computed using bins; n= number of regions analyzed where 1 region is 100 bins. p-value calculated by Wilcoxon testing.

Diifferential Gene Expression: Microarray EPC1-PHF1 Vs Mock K562

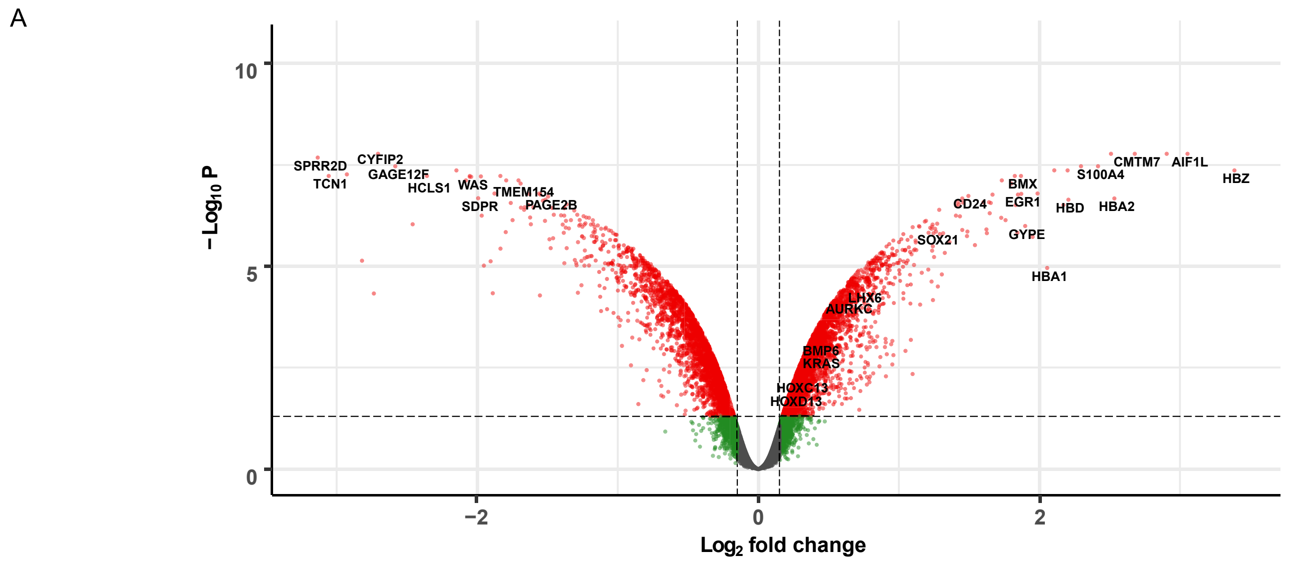


Genes Expression Changes at EPC1-PHF1 bound genes: Microarray correlation with FLAG ChIP-seq

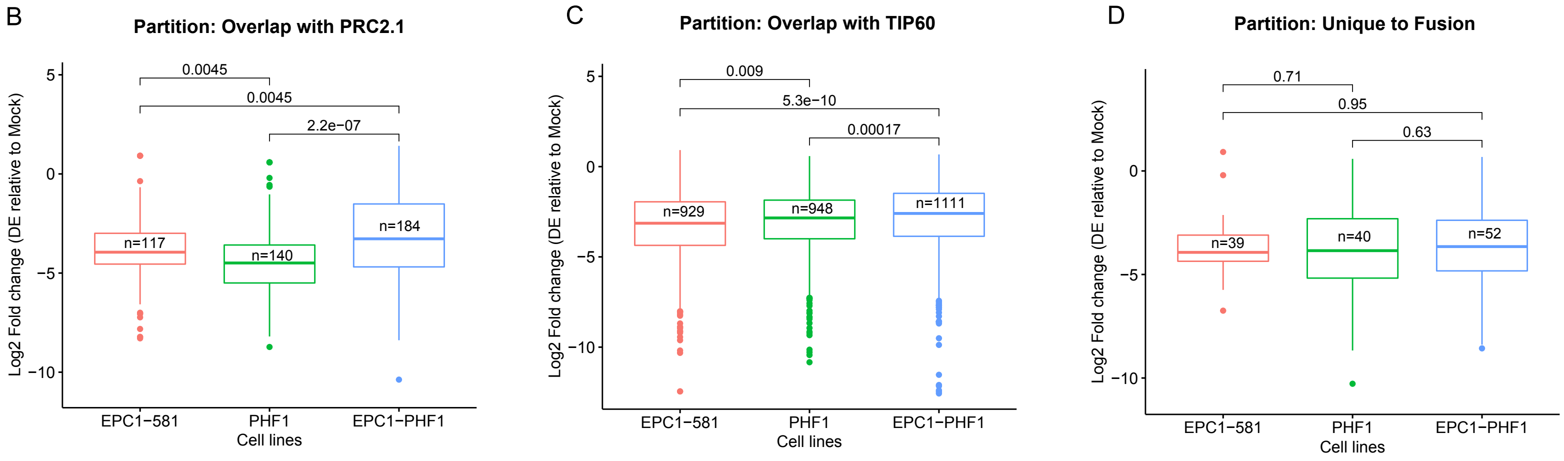

**<u>Figure S4</u> (Related to Figure 2): Gene transcription changes induced by EPC1-PHF1.**

**(A)** Volcano plot of differential gene expression analysis (microarray) comparing the EPC1-PHF1 expressing K562 cell line versus a Mock/empty vector K562 cell line.

**(B,C,D)** Gene expression changes at genes bound by EPC1-PHF1 in the isogenic cell lines and the indicated partitions. p-value calculated by Wilcoxon testing.

Sudarshan_Supplemental_FigS5

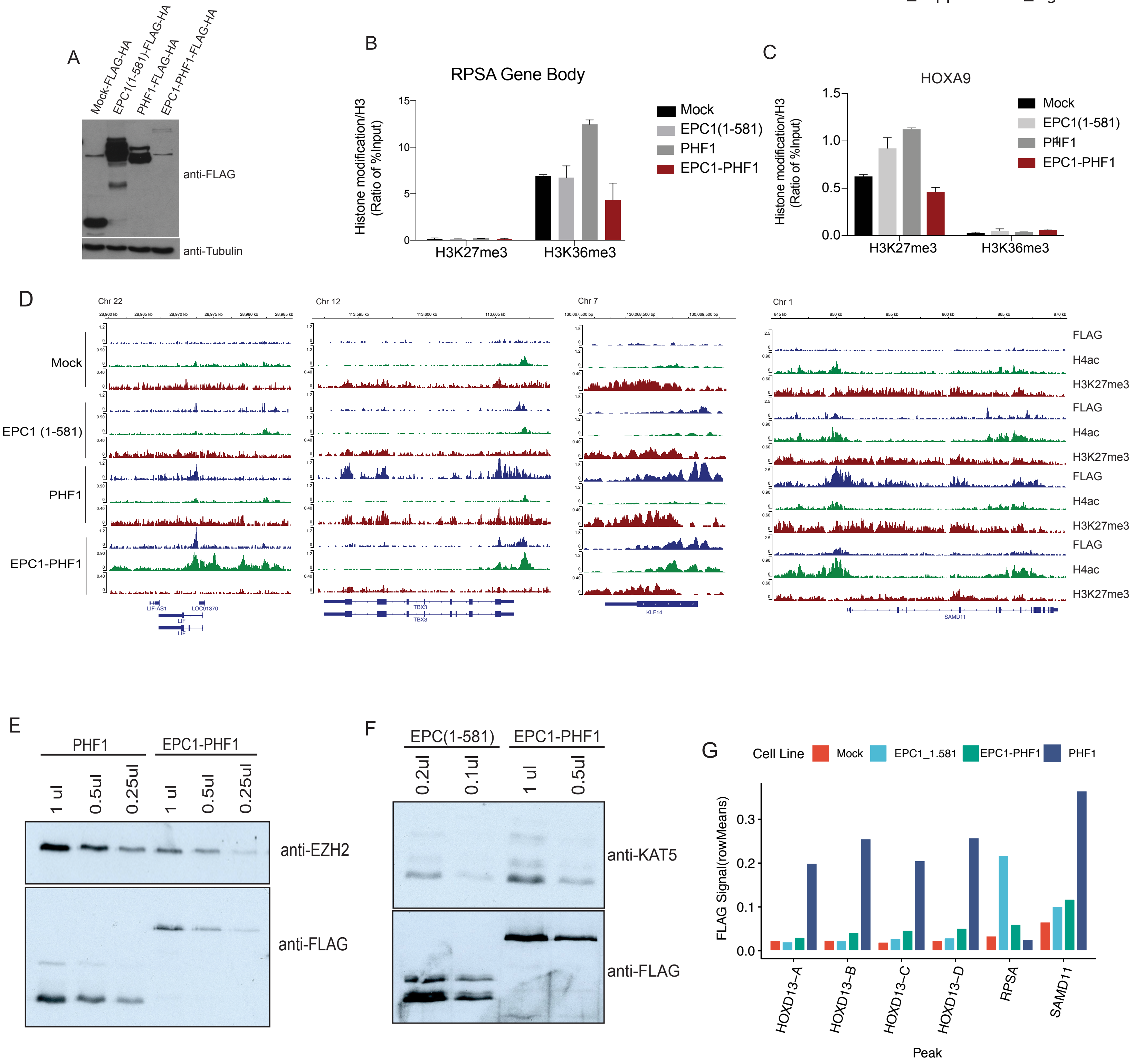

**<u>Figure S5</u> (Related to Figure 3):**

**(A)** Relative protein expression levels of EPC1(1-581), PHF1 and EPC1-PHF1 K562 cell lines used for ChIP-seq (in Figures 2, 3). EPC1-PHF1 is less expressed than other constructs. Western blotting from whole cell extracts was done with anti-FLAG and Tubulin is used as loading control.

**(B,C)** H3K27me3 and H3K36me3 levels at control gene regions: (B) ChlP-qPCR at a H3K36me3 positive locus, *RPSA* gene body; (C) ChlP-qPCR at a H3K27me3 positive region, *HOXA9*. (Values are represented as ratio of %IP/input chromatin of H3K36me3 or H3K27me3 and H3) (n=2, error bars represent Range of the values).

**(D)** ChIP-sequencing tracks of selected EPC1-PHF1 bound genes (LIF, TBX3, KLF14, SAMD11) showing increase in H4 acetylation and decrease in H3K27me3.

**(E,F)** Western blotting of purified fractions for normalization: (E) PHF1 and EPC1-PHF1 complexes were normalized for Histone Methyltransferase Assay (HMT) in Figure 3H using EZH2 antibody (5ul of Mock, 5ul of EPC1-PHF1 and 1.875ul of PHF1 were used for the assay). (F) EPC1(1-581) and EPC1-PHF1 complexes were normalized for Histone Acetyltransferase Assay (HAT) in Figure 3I using KAT5 antibody. (2ul of Mock, 2ul of EPC1-PHF1, 0.635ul of EPC1(1-581) were used for the assay)

**(G)** Histogram showing increased sequencing read means of EPC1-PHF1-FLAG ChIP signal versus mock control at the peaks of *de novo* acetylation detected is cells expressing the fusion near *HOXD13*. SAMD11 is a positive control for PHF1-FLAG reads while RPSA shows enrichment for EPC1(1-581)-FLAG reads.

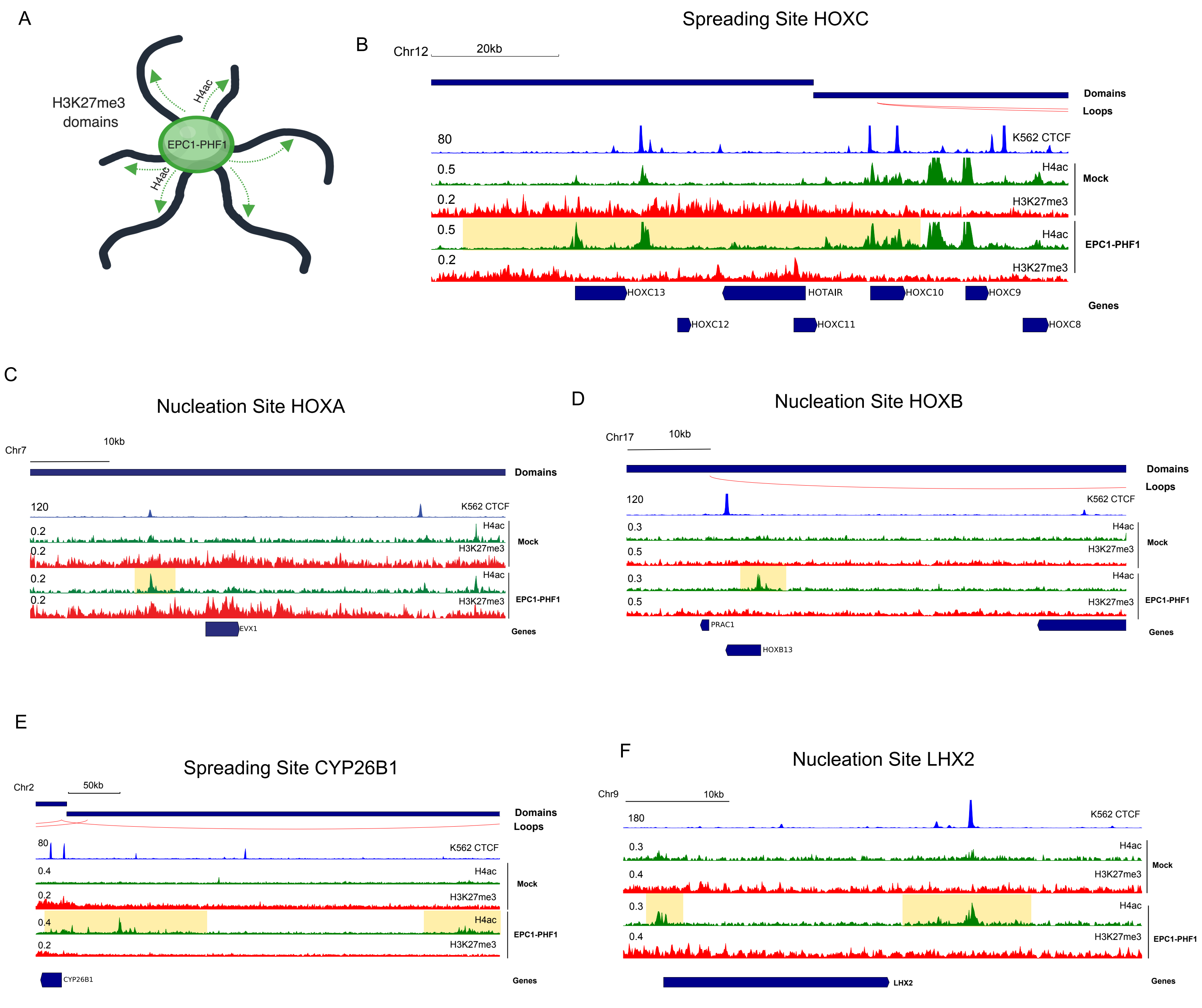
A
H3K27me3
domains
EPC1-PHF1
H4ac
H4ac
Spreading Site HOXC
B
Chr12
20kb
Domains
Loops
80
K562 CTCF
0.5
H4ac
Mock
0.2
H3K27me3
0.5
H4ac
EPC1-PHF1
0.2
H3K27me3
HOXC13
HOTAIR
HOXC10
HOXC9
Genes
HOXC12
HOXC11
HOXC8
C
Nucleation Site HOXA
Chr7
10kb
Domains
120
K562 CTCF
0.2
H4ac
Mock
0.2
H3K27me3
0.2
H4ac
EPC1-PHF1
0.2
H3K27me3
EVX1
Genes
D
Nucleation Site HOXB
Chr17
10kb
Domains
Loops
120
K562 CTCF
0.3
H4ac
Mock
0.5
H3K27me3
0.3
H4ac
EPC1-PHF1
0.5
H3K27me3
PRAC1
Genes
HOXB13
E
Spreading Site CYP26B1
Chr2
50kb
Domains
Loops
80
K562 CTCF
0.4
H4ac
Mock
0.2
H3K27me3
0.4
H4ac
EPC1-PHF1
0.2
H3K27me3
CYP26B1
Genes
F
Nucleation Site LHX2
Chr9
10kb
180
K562 CTCF
0.3
H4ac
Mock
0.4
H3K27me3
0.3
H4ac
EPC1-PHF1
0.4
H3K27me3
LHX2
Genes

<u>Figure S6</u> **(Related to Figure 3): Spreading of the EPC1-PHF1 complex and its associated histone marks.**

**(A)** Schematic representation of the possible mechanism of spreading of the EPC1-PHF1 complex and its associated histone marks, based on the reported mechanism of H3K27me3 spreading in cis and far cis through long range chromatin contacts (Oksuz et al., 2018).

**(B)** Representative ChIP-seq tracks at a spreading site - *HOXC* gene cluster, highlighted regions show changes in chromatin marks compared to control.

**(C)** Representative ChIP-seq tracks at a nucleation site - *EVX1* at *HOXA* gene cluster, highlighted regions show changes in chromatin marks compared to control.

**(D)** Representative ChIP-seq tracks at a nucleation site - *HOXB* gene cluster, highlighted regions show changes in chromatin marks compared to control.

**(E)** Representative ChIP-seq tracks at a spreading site - *CYP26B1*, proximal to *HOXD* gene cluster, highlighted regions show changes in chromatin marks compared to control.

**(F)** Representative ChIP-seq tracks at a nucleation site - *LHX2* (Lim Homeobox 2), highlighted regions show changes in chromatin marks compared to control.

A

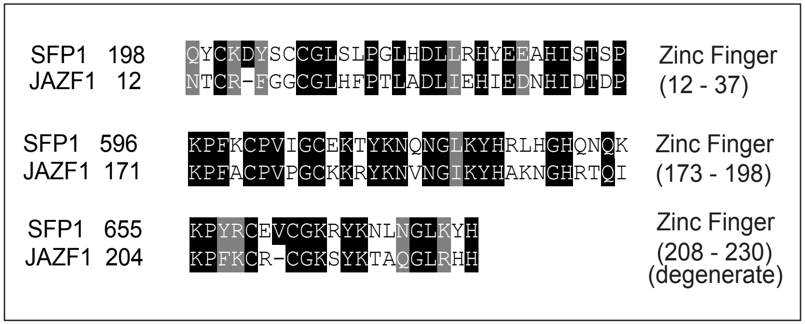


B

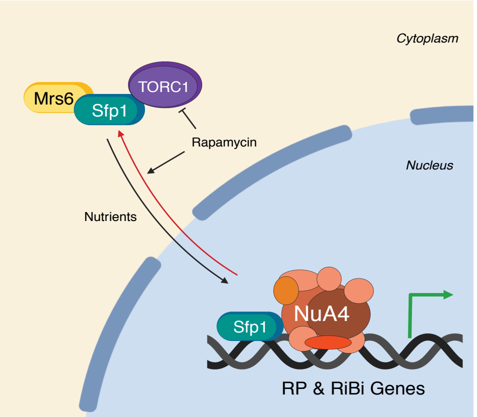


C

| PROTEIN | SPECTRAL COUNTS |
|---|---|
| TRRAP | 207 |
| EP400 | 127 |
| RUVBL2 | 48 |
| RUVBL1 | 44 |
| KAT5 | 17 |
| EPC2 | 16 |
| DMAP1 | 16 |
| MBTD1 | 17 |
| EPC1 | 11 |
| JAZF1 | 10 |
| BAF53a | 24 |
| ING3 | 3 |
| BRD8 | 2 |

D

| | PROTEIN | SPECTRAL COUNTS |
|---|---|---|
| TIP60 COMPLEX | TRRAP | 1071 |
| | EP400 | 595 |
| | RUVBL1 | 352 |
| | RUVBL2 | 346 |
| | EPC2 | 231 |
| | BRD8 | 186 |
| | EPC1 | 155 |
| | KAT5 | 143 |
| | ING3 | 118 |
| | DMAP1 | 117 |
| | ACTB | 107 |
| | BAF53a | 94 |
| | MBTD1 | 94 |
| | MRG15 | 70 |
| | MRGX | 56 |
| | VPS72 | 54 |
| | YEATS4 | 51 |
| | MEAF6 | 43 |
| | MRGBP | 32 |
| | JAZF1 | 33 |
| | NR2C2 | 34 |
| | NR2C1 | 15 |
| | H2B | 09 |
| | H2A.Z | 05 |

E

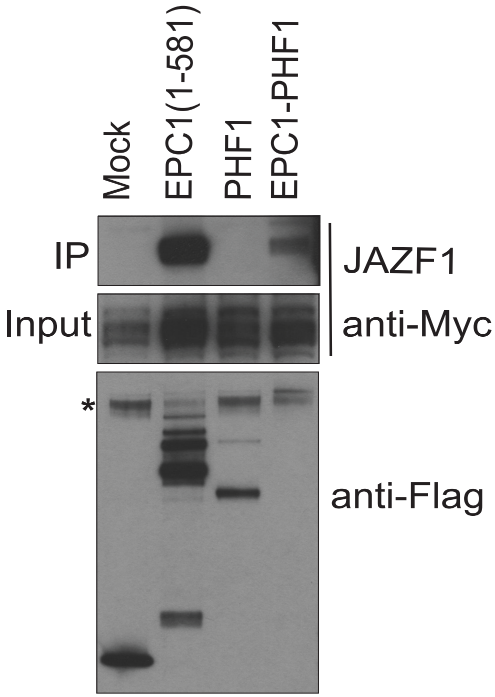

**Figure S7 (Related to Figure 4): JAZF1 associates with the NuA4/TIP60 complex.**

**(A)** Protein sequence alignment: Human JAZF1 and yeast Sfp1 show homology in the zinc-finger domains. Black boxes show exact amino acid conservation, grey boxes show conservation of amino acids with similar polarity.

**(B)** Model depicting the relationship between SFP1 and NuA4 in *S. cerevisiae*. Sfp1 is a transcriptional activator that responds to nutrients and stress, being regulated by TORC1 and Mrs6. In the nucleus, Sfp1 and the NuA4 complex co-operate to regulate the expression of Ribosomal Protein (RP) and Ribosomal Biogenesis (RiBi) genes.

**(C)** JAZF1 interacts with the TIP60 complex. Table of proteins (numbers represent total spectral counts) co-purified with JAZF1 in the chromatin fractions from HEK293 cells upon multiple cell compartment affinity purification coupled to tandem mass spectrometry analysis (MCC-AP-MS/MS) (Data from two biological replicates; Interactions with an FDR below 20% in both replicate runs and interactions observed in only one replicate run with an FDR < 10% are reported).

**(D)** Mass Spectrometry analysis of the JAZF1-3xFLAG-2xStrep affinity purified from K562 cells. Table with total spectral counts of peptides present in the tandem affinity purified fraction shown in Figure 4A-B. Peptides highlighted in purple are NuA4/TIP60 complex components, peptides highlighted in red are other known JAZF1 interacting proteins, in green are histones.

**(E)** Immunoprecipitation of JAZF1 with the EPC1-PHF1 fusion. Myc-tagged JAZF1 construct was transiently expressed in FLAG-tagged K562 cell lines expressing Tag (Mock), EPC1(1-581), PHF1 or EPC1-PHF1) from the *AAVS1* locus. Immunoprecipitation was performed with Anti-FLAG-resin and analyzed by western blotting with Anti-Myc antibody. JAZF1 interacts with EPC1(1-581) and EPC1-PHF1. See Figure 4C for data with JAZF1(1-124) construct.

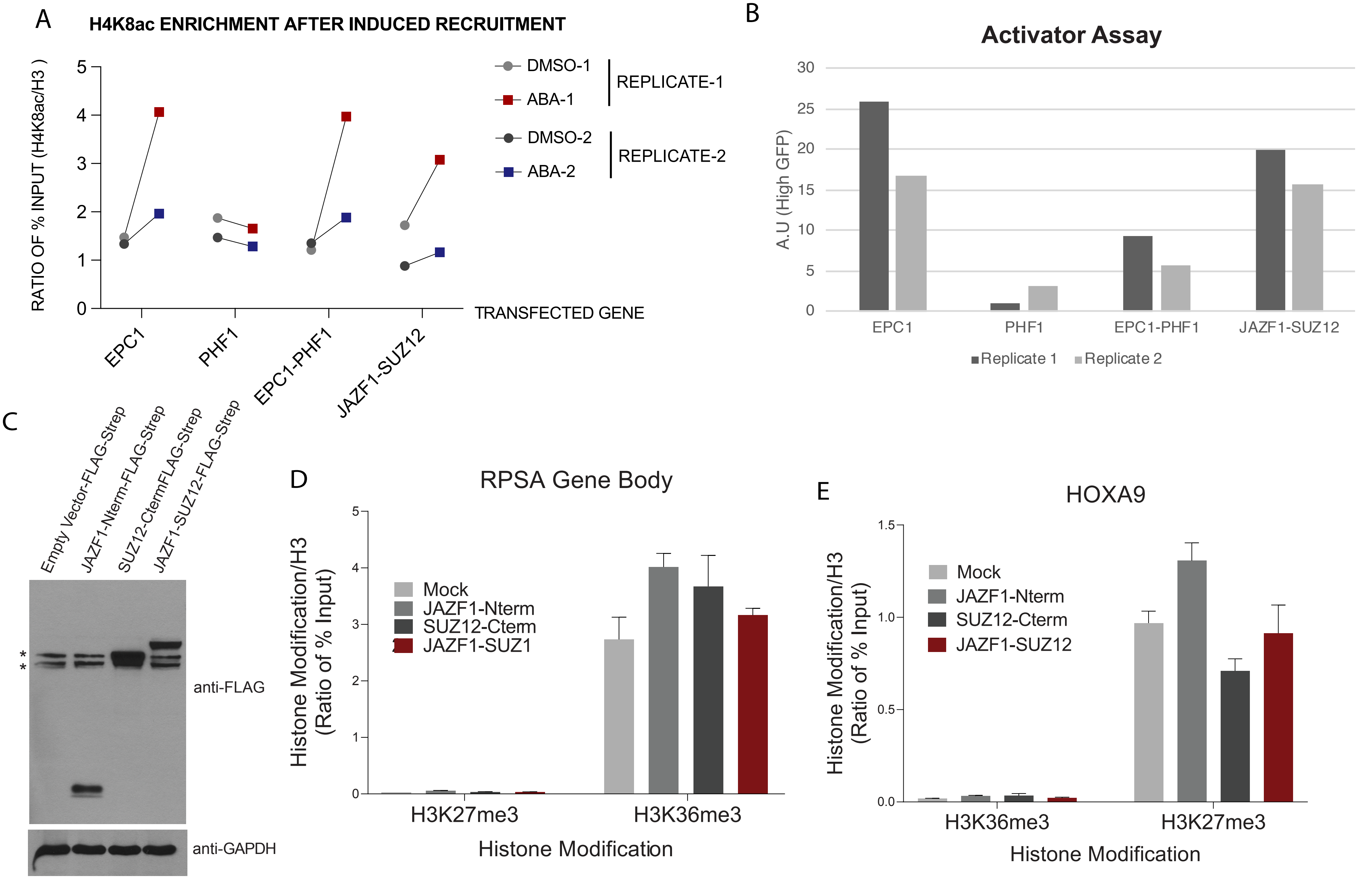
A
H4K8ac ENRICHMENT AFTER INDUCED RECRUITMENT
RATIO OF % INPUT (H4K8ac/H3 )
0
1
2
3
4
5
DMSO-1
ABA-1
REPLICATE-1
DMSO-2
ABA-2
REPLICATE-2
TRANSFECTED GENE
EPC1
PHF1
EPC1-PHF1
JAZF1-SUZ12
B
Activator Assay
A.U (High GFP)
0
5
10
15
20
25
30
EPC1
PHF1
EPC1-PHF1
JAZF1-SUZ12
Replicate 1
Replicate 2
C
Empty Vector-FLAG-Strep
JAZF1-Nterm-FLAG-Strep
SUZ12-CtermFLAG-Strep
JAZF1-SUZ12-FLAG-Strep
*
*
anti-FLAG
anti-GAPDH
D
RPSA Gene Body
Histone Modification/H3
(Ratio of % Input)
0
1
2
3
4
5
Mock
JAZF1-Nterm
SUZ12-Cterm
JAZF1-SUZ1
H3K27me3
H3K36me3
Histone Modification
E
HOXA9
Histone Modification/H3
(Ratio of % Input)
0.0
0.5
1.0
1.5
Mock
JAZF1-Nterm
SUZ12-Cterm
JAZF1-SUZ12
H3K36me3
H3K27me3
Histone Modification

**<u>Figure S8</u> (Related to Figure 5): The JAZF1-SUZ12 fusion mislocalizes histone marks.**

**(A)** H4K8ac ChIP-qPCR at EGFP reporter (+300bp after promoter) after induced recruitment of indicated proteins.

**(B)** Flow cytometry analysis of replicates used for ChIP-qPCR in (A). The histogram shows % of cells transfected with EPC1, PHF1, EPC1-PHF1 or EPC1 (1-581) expressing high GFP upon ABA treatment.

**(C)** Protein expression level of JAZF1-SUZ12, JAZF1-Nterm, SUZ12-C-term and empty tag (3xFLAG-2xStrep) from the *AAVS1* locus in K562 cells. Anti-FLAG western blotting on whole cell extracts and GAPDH is used as loading control.

**(D, E)** H3K27me3 and H3K36me3 levels in JAZF1-SUZ12 expressing cells at control loci. (D) ChlP-qPCR at a H3K36me3 positive locus, *RPSA* gene body; (E) ChlP-qPCR at a H3K27me3 positive region, *HOXA9*. (Values are represented as ratio of % of input chromatin of H3K36me3 or H3K27me3 and H3) (n=2, error bars represent Range of the values).

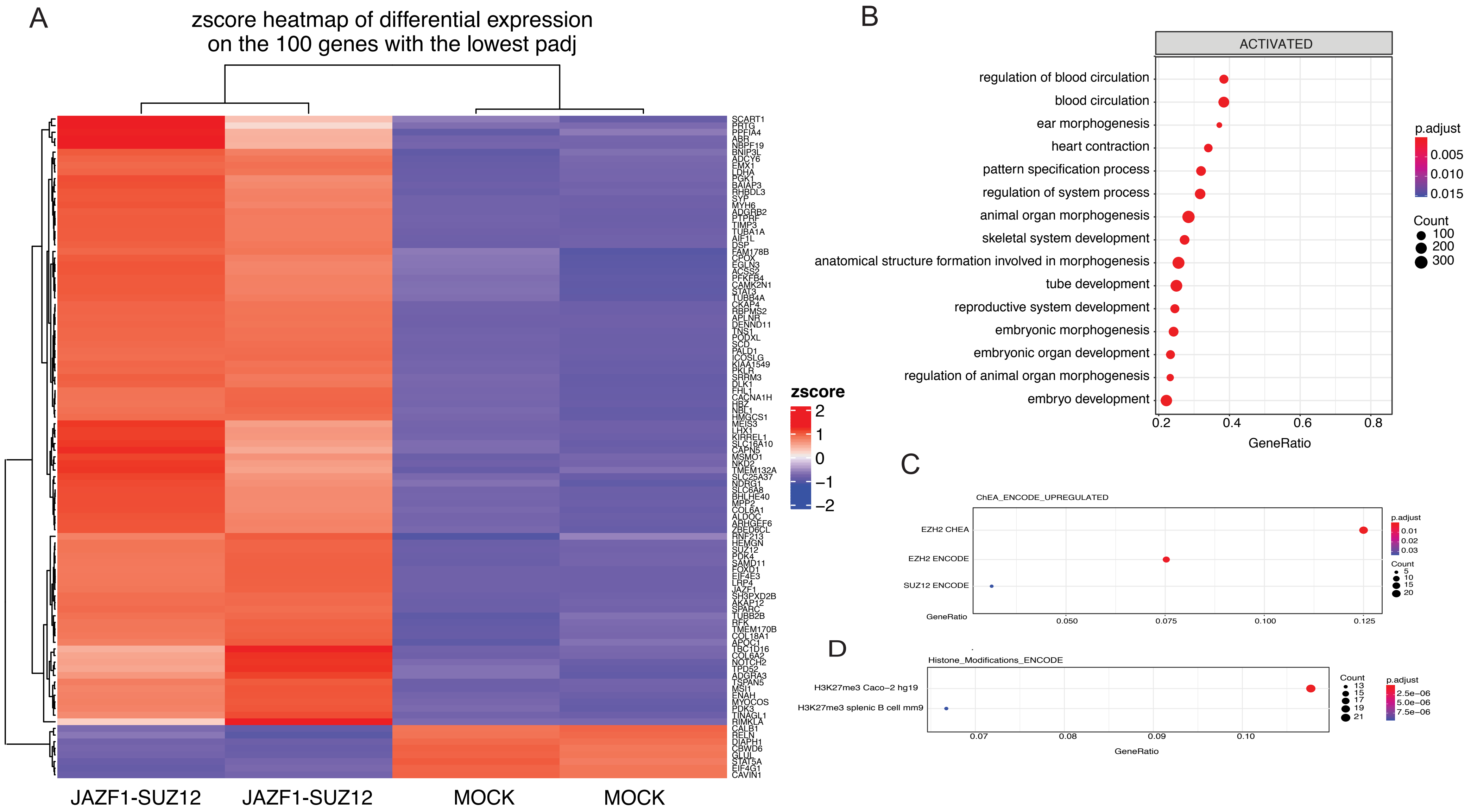
A
zscore heatmap of differential expression
on the 100 genes with the lowest padj
SCART1
PRTG
PPFIA4
ABR
NBPF19
BNIP3L
ADCY6
EMX1
LDHA
PGK1
BAIAP3
RHBDL3
SYP
MYH6
ADGRB2
PTPRF
TIMP3
TUBA1A
AIF1L
DSP
FAM178B
CPOX
EGLN3
ACSS2
PFKFB4
CAMK2N1
STAT3
TUBB4A
CKAP4
RBPMS2
APLNR
DENND11
TNS1
PODXL
SCD
PALD1
ICOSLG
KIAA1549
PKLR
SRRM3
DLK1
FHL1
CACNA1H
HBZ
NBL1
HMGCS1
MEIS3
LHX1
KIRREL1
SLC16A10
CAPN5
MSMO1
NKD2
TMEM132A
SLC25A37
NDRG1
SLC6A8
BHLHE40
MPP2
COL6A1
ALDOC
ARHGEF6
ZBED6CL
RNF213
HEMGN
SUZ12
PDK4
SAMD11
FOXD1
EIF4E3
LRP4
JAZF1
SH3PXD2B
AKAP12
SPARC
TUBB2B
RFK
TMEM170B
COL18A1
APOC1
TBC1D16
COL6A2
NOTCH2
TPD52
ADGRA3
TSPAN5
MSI1
ENAH
MYOCOS
PDK3
TINAGL1
RIMKLA
CALB1
RELN
DIAPH1
CBWD6
GLUL
STAT5A
EIF4G1
CAVIN1
zscore
2
1
0
−1
−2
JAZF1-SUZ12
JAZF1-SUZ12
MOCK
MOCK
B
ACTIVATED
regulation of blood circulation
blood circulation
ear morphogenesis
heart contraction
pattern specification process
regulation of system process
animal organ morphogenesis
skeletal system development
anatomical structure formation involved in morphogenesis
tube development
reproductive system development
embryonic morphogenesis
embryonic organ development
regulation of animal organ morphogenesis
embryo development
0.2
0.4
0.6
0.8
GeneRatio
p.adjust
0.005
0.010
0.015
Count
100
200
300
C
ChEA_ENCODE_UPREGULATED
EZH2 CHEA
EZH2 ENCODE
SUZ12 ENCODE
GeneRatio
0.050
0.075
0.100
0.125
p.adjust
0.01
0.02
0.03
Count
5
10
15
20
D
Histone_Modifications_ENCODE
H3K27me3 Caco−2 hg19
H3K27me3 splenic B cell mm9
0.07
0.08
0.09
0.10
GeneRatio
Count
13
15
17
19
21
p.adjust
2.5e−06
5.0e−06
7.5e−06

**Figure S9 (Related to Figure 5): Gene expression changes in JAZF1-SUZ12 expressing cells**

**(A)** Heatmap of differential expression analysis comparing JAZF1-SUZ12 expressing K562 cell line and empty vector K562 cell line (Mock).

**(B)** GSEA using the Gene Ontology database (Biological Processes) was performed on upregulated genes in JAZF1-SUZ12 expressing K562 cell line. 15 associated significant gene sets with lowest padj values (< 0.05) calculated by FDR method are presented.

**(C)** Pathway over-representation analysis using the ENCODE and ChEA Consensus TFs from ChIP-X library was performed on upregulated genes in JAZF1-SUZ12 expressing K562 cells. The associated Transcription factors (EZH2 and SUZ12) with the padj values below 0.05 are presented.

**(D)** Pathway over-representation analysis using the ENCODE Histone Modifications database (2015) was performed on upregulated genes in JAZF1-SUZ12 expressing K562 cells. The associated histone modification (H3K27me3) with the padj values below 0.05 are presented.

Sudarshan_Supplemental_FigS10

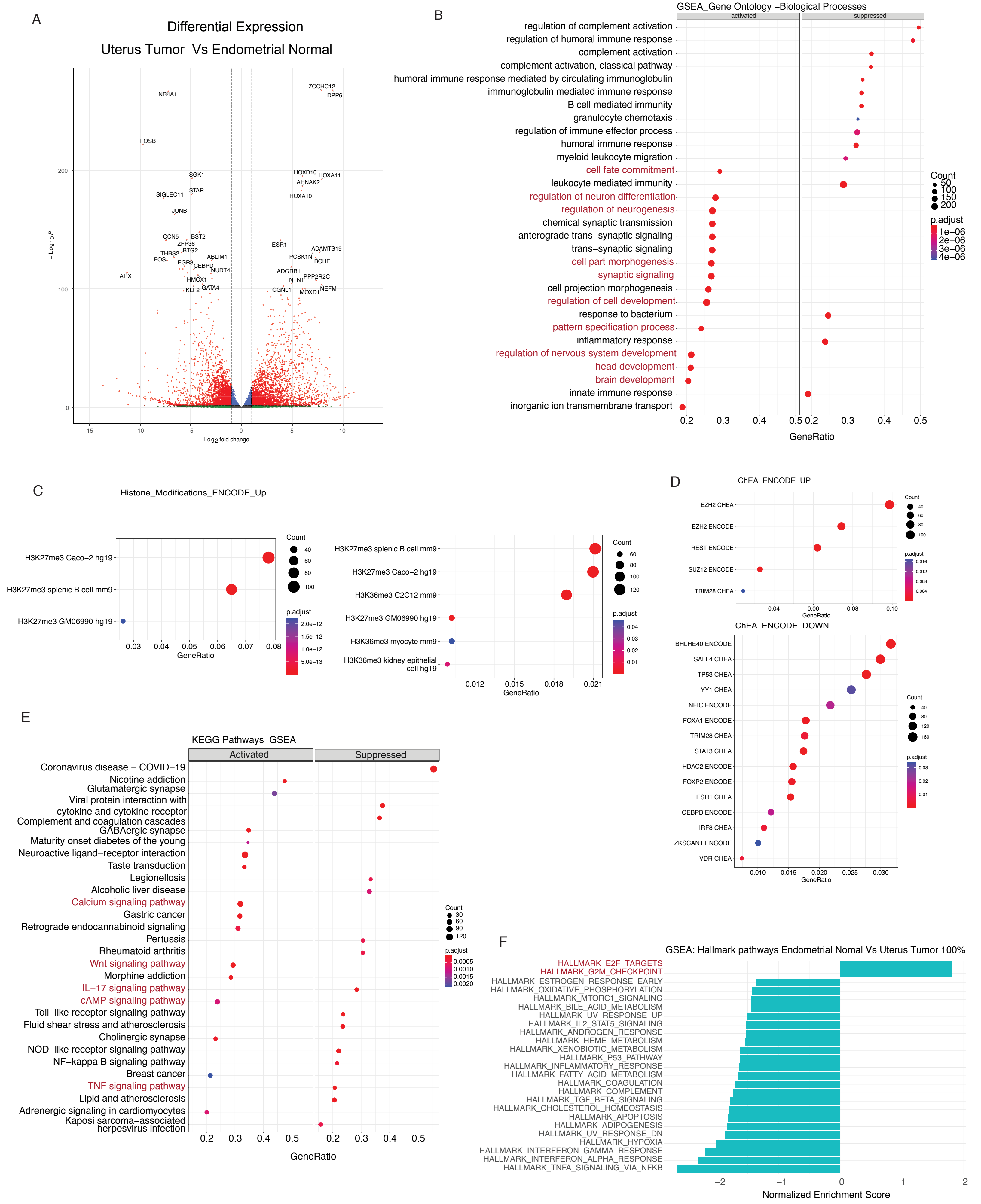

<u>Figure S10</u> **(Related to Figure 5): Differential Expression analysis in LG-ESS patient samples showing the presence of JAZF1-SUZ12 fusion**

**(A)** Volcano plot showing significantly differentially expressed genes from pairwise comparisons of Uterus Tumor to Endometrial Normal tissue.

**(B,E,F)** GSEA using the Hallmark Gene Sets from Gene Ontology – Biological pathways, KEGG Pathways and Molecular Signatures Database (MSigDB) was performed on differentially expressed genes in Uterus Tumor patient sample compared to the Endometrial normal tissue. Associated significant gene sets with lowest padj values (< 0.05) calculated by FDR method are presented.

**(C)** Pathway over-representation analysis using the ENCODE Histone Modifications database (2015) was performed on differentially expressed genes in Uterus Tumor sample in comparison to the Endometrial normal tissue. The associated histone modifications with the padj values (hypergeometric distribution) below 0.05 are presented.

**(D)** Pathway over-representation analysis using the ENCODE and ChEA Consensus TFs from ChIP-X library was performed on differentially expressed genes in Uterus Tumor sample in comparison to the Endometrial normal tissue. The associated Transcription factors with the padj values (hypergeometric distribution) below 0.05 are presented.

Sudarshan_Supplemental_FigS11

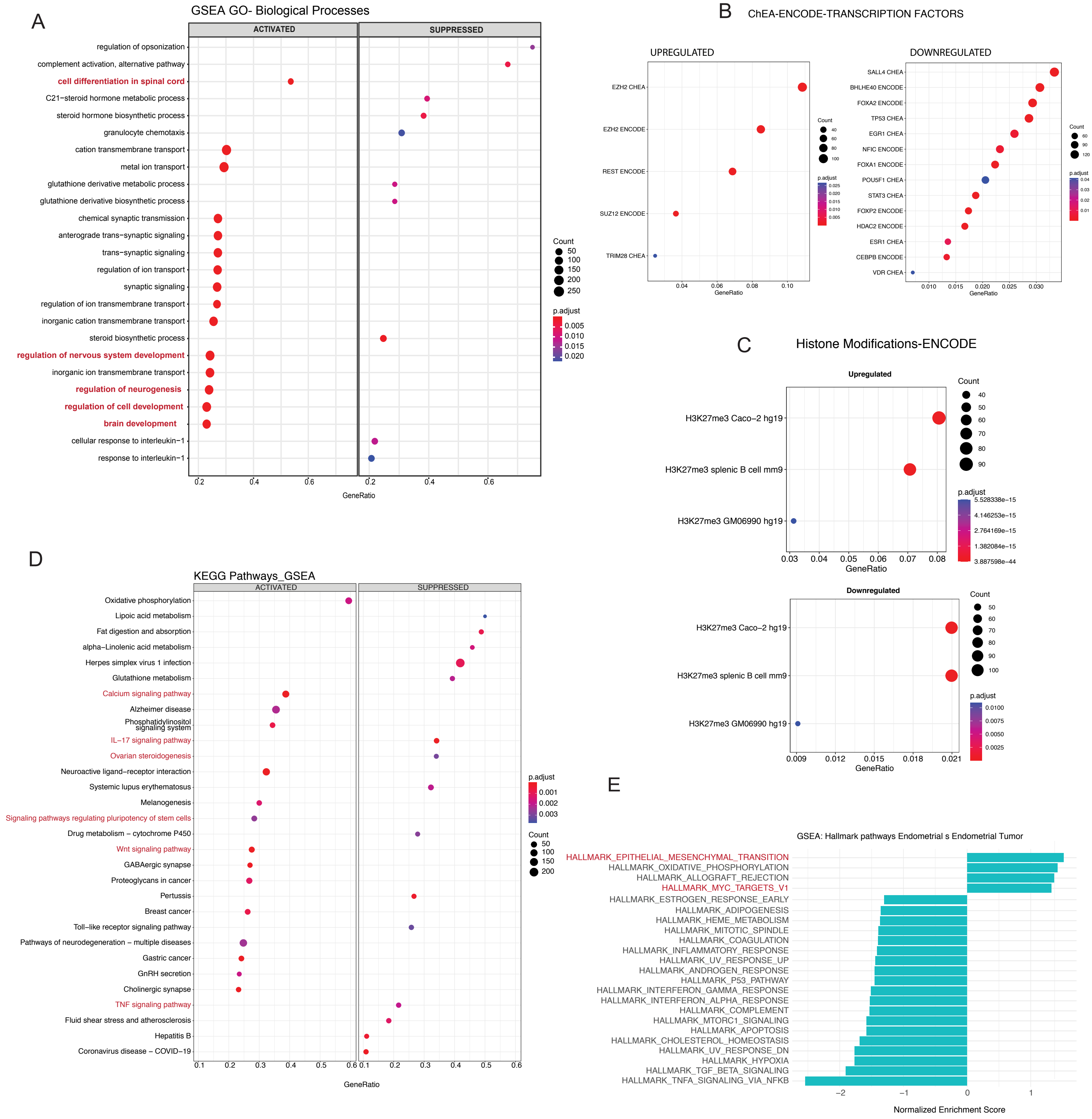

<u>Figure S11</u> (**Related to Figure 5): Differential Expression Pathway analysis in LG-ESS patient samples showing the presence of JAZF1-SUZ12 fusion**

**(A)** GSEA using the Gene Ontology database (Biological Processes) was performed on differentially expressed genes in Endometrial Tumor sample in comparison to the paired Endometrial Normal tissue from the same patient. 25 of the associated significant gene sets with lowest padj values (< 0.05) calculated by FDR method are presented.

**(B)** Pathway over-representation analysis using the ENCODE and ChEA Consensus TFs from ChIP-X library was performed on differentially expressed genes in Endometrial Tumor sample in comparison to the paired Endometrial normal tissue. The associated Transcription factors with the padj values (hypergeometric distribution) below 0.05 are presented.

**(C)** Pathway over-representation analysis using the ENCODE Histone Modifications database (2015) was performed on differentially expressed genes in Endometrial Tumor sample in comparison to the paired Endometrial normal tissue. The associated histone modifications with the padj values (hypergeometric distribution) below 0.05 are presented.

**(D,E)** GSEA using the KEGG Pathways and Molecular Signatures Database (MSigDB) was performed on differentially expressed genes in Endometrial Tumor sample in comparison to the paired Endometrial Normal tissue. Associated significant gene sets with lowest padj values (< 0.05) calculated by FDR method are presented.

### SUPPLEMENTARY METHODS

#### Acid Extraction of Histones

EPC1-PHF1-Flag Tagged K562 and control cells were grown to $10^7$ cell density in 100mm dishes, harvested, washed twice in 1XPBS. Cells were pelleted and resuspended in 1ml lysis buffer (PBS containing 0.5% Triton X 100 (v/v), 2 mM phenylmethylsulfonyl fluoride (PMSF), 0.02% (w/v) NaN3, 5mM sodium Butyrate) for 10 min, nuclei were spun down, resuspended in half volume of lysis buffer and centrifuged. Pellet was resuspended in 250 µl 0.2N HCl, incubated overnight at 4C, centrifuged. Supernatant containing acid extracted histones was neutralized with 1/10 volume of NaOH. Histones were quantified by Bradford assay, resolved on 15% SDS-PAGE, and Immunoblotted with anti-H3K27me3 and anti-H4 antibodies. **(**https://www.abcam.com/protocols/histone-extraction-protocol-for-western-blot**)**

**Colony formation assay:** HEK293T were transfected using the calcium phosphate method and then grown under selection with 300ug/ml hygromycin B for two weeks. Colonies that formed were fixed with methanol and colored with a solution of Giemsa stain for easier visualization. Plates were scanned using an Epson Perfection 4870 scanner; colonies in the images were counted using Alpha Imager software.

**Multiple cell compartment affinity purification coupled to tandem mass spectrometry (MCC-AP-MS/MS)** : Inducible EcR-293 stable cell line containing pMZI empty vector as control, pMZI-JAZF1 were generated as previously described (Jeronimo et al. 2007). Protein expression was induced with 3 µM ponasterone A (Life Technologies) and cells were harvested after 48h. Protein-

protein interactions of JAZF1 were identified using two biological replicate runs for each bait of the multiple cell compartment affinity purification coupled to tandem mass spectrometry (MCC-AP-MS/MS) procedure, as recently described (Lavallee-Adam et al. 2013), (Cloutier et al. 2014)). The eluates were precipitated with trichloroacetic acid (TCA) and stored at -80 °C until liquid chromatography (LC) MS/MS analysis. Protein were solubilized, digested with trypsin and analysed by tandem mass spectrometry as previously described (Cloutier et al. 2014). Protein database searching was performed with Mascot 2.2 (Matrix Science) against the human NCBInr protein database. Reliability assessment of protein-protein interactions identified with MCC-AP-MS/MS was performed using our previously published software package Decontaminator (Lavallee-Adam et al. 2011). pMZI empty vector controls were used to train Decontaminator's Bayesian inference algorithm. Decontaminator assigned a False Discovery Rate (FDR) to each protein-protein interaction in our dataset using a leave-one-out procedure. Interactions with a FDR below 20% in both replicate runs of MCC-AP-MS/MS and interactions observed in only one replicate run with a FDR < 10% were reported.

**Mass spectrometry Analysis of Purified Complexes :** Purified complexes were run on preparative gels (Bolt 12% Bis-Tris, Invitrogen) over 1 cm, to stack all proteins into 1 band, stained with Sypro Ruby (Bio-Rad) and excised under UV lamp. The gel slice was reduced, alkylated, and digested based on the original protocol (Wilm et al. 1996). Briefly, proteins were reduced with 10 mM DTT at 56°C for 15 minutes and alkylated with 100 mM iodoacetamide for 15 minutes. Trypsin digestion was performed using modified porcine trypsin (Sequencing grade, Sigma) at 37°C for 16 h. Digestion products were extracted sequentially using 25 mM ammonium bicarbonate, then

50% acetonitrile with 5% formic acid and finally 100% acetonitrile. The recovered extracts were pooled, vacuum centrifuge dried and stored at -80°C until MS analysis.

**Proteins identification by mass spectrometry – TripleTOF 5600:** Each sample (5 µL) was directly loaded at 400 nL/min onto an equilibrated HPLC column. The peptides were eluted from the column over a 90 min gradient generated by a NanoLC-Ultra 1D plus (Eksigent, Dublin CA) nano-pump and analysed on a TripleTOF 5600 instrument (AB SCIEX, Concord, Ontario, Canada). The gradient was delivered at 200 nL/min starting from 2% acetonitrile with 0.1% formic acid to 35% acetonitrile with 0.1% formic acid over 90 min followed by a 15 min clean-up at 80% acetonitrile with 0.1% formic acid, and a 15 min equilibration period back to 2% acetonitrile with 0.1% formic acid, for a total of 120 min. To minimize carryover between each sample, the analytical column was washed for 3 h by running an alternating sawtooth gradient from 35% acetonitrile with 0.1% formic acid to 80% acetonitrile with 0.1% formic acid, holding each gradient concentration for 5 min. Analytical column and instrument performance were verified after each sample by loading 30 fmol bovine serum albumin (BSA) tryptic peptide standard (Michrom Bioresources Inc. Fremont, CA) with 60 fmol α-casein tryptic digest and running a short 30 min gradient. TOF MS calibration was performed on BSA reference ions before running the next sample to adjust for mass drift and verify peak intensity. The instrument method was set to data dependent acquisition (DDA) mode, which consisted of one 250 milliseconds (ms) MS1 TOF survey scan from 400–1300 Da followed by 20 100 ms MS2 candidate ion scans from 100–2000 Da in high sensitivity mode. Only ions with a charge of 2+ to 4+ that exceeded a threshold of 200 cps were selected for MS2, and former precursors were excluded for 10 s after one occurrence.

**Proteins identification by mass spectrometry – Orbitrap Fusion:** Peptide samples were separated by online reversed-phase (RP) nanoscale capillary liquid chromatography (nanoLC) and analyzed by electrospray mass spectrometry (ESI MS/MS). The experiments were performed with a Dionex UltiMate 3000 nanoRSLC chromatography system (Thermo Fisher Scientific) connected to an Orbitrap Fusion mass spectrometer (Thermo Fisher Scientific) equipped with a nanoelectrospray ion source. Peptides were trapped at 20 µl/min in loading solvent (2% acetonitrile, 0.05% TFA) on a 5mm x 300 µm C18 pepmap cartridge pre-column (Thermo Fisher Scientific) during 5 minutes. Then, the pre-column was switch online with a self-made 50 cm x 75 um internal diameter separation column packed with ReproSil-Pur C18-AQ 3-µm resin (Dr. Maisch HPLC) and the peptides were eluted with a linear gradient from 5-40% solvent B (A: 0,1% formic acid, B: 80% acetonitrile, 0.1% formic acid) in 90 minutes, at 300 nL/min. Mass spectra were acquired using a data dependent acquisition mode using Thermo XCalibur software version 3.0.63. Full scan mass spectra (350 to 1800m/z) were acquired in the orbitrap using an AGC target of 4e5, a maximum injection time of 50 ms and a resolution of 120 000. Internal calibration using lock mass on the m/z 445.12003 siloxane ion was used. Each MS scan was followed by acquisition of fragmentation spectra of the most intense ions for a total cycle time of 3 seconds (top speed mode). The selected ions were isolated using the quadrupole analyzer in a window of 1.6 m/z and fragmented by Higher energy Collision-induced Dissociation (HCD) with 35% of collision energy. The resulting fragments were detected by the linear ion trap in rapid scan rate with an AGC target of 1E4 and a maximum injection time of 50ms. Dynamic exclusion of previously fragmented peptides was set for a period of 20 sec and a tolerance of 10 ppm.

Mass spectrometry data were stored, searched, and analyzed using the ProHits laboratory information management system (LIMS) platform (Liu et al. 2016). Within ProHits, AB SCIEXWIFF files were first converted to an MGF formatusing WIFF2MGF converter and to an mzML format using ProteoWizard (v3.0.4468) (Kessner et al. 2008) and the AB SCIEX MS Data Converter (V1.3

beta). The mzML files were searched using Mascot (v2.3.02). The spectra were searched with the RefSeq database (version 57, January 30th, 2013) acquired from NCBI against a total of 72,482 human and adenovirus sequences supplemented with common contaminants from the Max Planck Institute (http://141.61.102.106:8080/share.cgi?ssid=0f2gfuB) and the Global Proteome Machine (GPM; http://www.thegpm.org/crap/index.html). The database parameters were set to search for tryptic cleavages, allowing up to two missed cleavage sites per peptide with a mass tolerance of 40 ppm for precursors with charges of 2+ to 4+ and a tolerance of +/- 0.15 amu for fragment ions. Deamidated asparagine and glutamine and oxidized methionine were allowed as variable modifications.

**SUPPLEMENTARY TABLES**

**Table S1: related to Figure 2 and 3 and Supplementary figure 2A, 2B, 3A**
Quality control FASTQC
annotated Peak Calls- FLAG, H4ac, 27me3
Annotated partitions of EPC1-PHF1 Flag peaks
Flag enrichment (Row means) at EPC1-PHF1 H4ac peaks
Microarray Differential Expression analysis
EPC1-PHF1-HA CUT and RUN peak calls
Antibodies used
ChiP-qPCR primers used

**Table S2: related to Figure 5 and Supplementary figure 5B**
JAZF1-SUZ12- k562 -RNA-seq-differential expression analysis
JAZF1-SUZ12- k562 -RNA-seq-GO enrichment
JAZF1-SUZ12- k562 -RNA-seq- ChEA-ENCODE TF enrichment
JAZF1-SUZ12- k562 -RNA-seq- Encode Histone Mark enrichment

**Table S3: related to Figure 5 and Supplementary figure 5C and 5D**
Star fusion prediction of fusions
ESS patient sample-RNA-seq-differential expression analysis, TPM, DE
Paired Endometrial cancer vs normal GSEA : GO , KEGG, Hallmark Pathways, ChEA-Encode TF, Encode Histone mark.
Uterus Tumor Vs Endometrial Normal GSEA: GO, KEGG, Hallmark, ChEA, histone mark
